# Fungal–beetle networks in deadwood are modular and shaped by tree species and deadwood type

**DOI:** 10.64898/2026.03.23.713619

**Authors:** Maria Faticov, Anders Dahlberg, Joakim Hjalten, Therese Lofroth, Anne-Maarit Hekkala

## Abstract

Deadwood is a key habitat for forest biodiversity, yet how tree species and deadwood type shape linked fungal–beetle communities remain poorly understood. We explored saproxylic fungi and beetles in a large-scale restoration experiment on birch, pine, and spruce deadwood created as burned standing trees, felled logs, girdled trees, high stumps, and uprooted trees. As expected, we found that tree species was the main driver of both fungal and beetle community composition, while deadwood type was the second most important driver. Fungal–beetle community correlations were context dependent: significant multivariate correlations were detected for pine and spruce, but not birch, and were strongest in burned standing pine, burned standing spruce, and girdled spruce. Across all tree species and deadwood types, fungal–beetle co-occurrence networks were consistently less nested and more modular than expected by chance, indicating structured, compartmentalized associations of fungi and beetles even within single deadwood units.

**Synthesis:** These results show that maintaining diverse tree species and deadwood types is essential to retain specialized multitrophic communities and the ecological processes they support.

## Introduction

Deadwood is a key structural and functional component of forests, serving as a hotspot of biodiversity and an essential part of nutrient cycling and decomposition (Stokland *et al*., 2012). As forestry typically removes trees for timber and pulp, the amount and quality of coarse deadwood that historically supported saproxylic diversity has been dramatically reduced and altered over the past century (Gibb *et al*., 2005; Stenbacka *et al*., 2010). Deadwood supports large proportion of forest biodiversity (Stokland *et al*., 2012), serving as a key substrate for decomposer species such as fungi and beetles, but also offer habitat for nitrogen-fixing bacteria (Tláskal & Baldrian, 2021) and, at later decay stages, other root and soil associated microorganisms (Purahong *et al*., 2022). Notably, fungi and beetles are the major saproxylic organisms, species rich and the ones in charge of the decomposition in deadwood (Siitonen 2001; Seibold *et al*., 2021). The enzymes of fungi decompose dead wood by breaking down cellulose and lignin, while beetles contribute to fragmentation, nutrient turnover, and fungal dispersal through vectoring of spores (Ulyshen, 2016). Together, these groups form interacting assemblages. To mitigate negative biodiversity impacts of forest management, preservation and creation of coarse dead wood is a key component in retention management approaches and restoration (Franklin *et al*., 1987; Gustafsson *et al*., 2010; Sandström *et al*., 2019). Yet the mechanisms by which different deadwood types affect multitrophic interactions remain poorly understood, making the outcome of restoration and retention effort for dead wood biodiversity and decomposition processes difficult to predict.

Interactions among fungi and saproxylic beetles in deadwood form complex ecological networks that are the bases of community assembly processes (Wende *et al*., 2017; Birkemoe *et al*., 2018; Jacobsen *et al*., 2018a). The organization of these multitrophic networks can largely be described by three key structural properties: nestedness, connectance and modularity. Nestedness reflects a hierarchical pattern in which specialists interact mainly with subsets of generalist species (Bascompte *et al*., 2003; Almeida-Neto *et al*., 2008). Such networks are typically robust, as generalist taxa acts as stabilizing nodes that buffer against local species extinctions and changes in the environment, partly because generalist species are often functionally redundant and can replace one another. Notably, the recent studies have demonstrated that beetle-fungus interactions in deadwood to be “anti-nested” with much lower nestedness than expected by chance (Jacobsen *et al*., 2018b). This means that interactions happen not among generalist species who interact with many other species withing the network but instead are more species-specific. Connectance, the proportion of realized to potential links between species, represents the overall density and complexity of interactions (Dunne *et al*., 2002; Gilbert, 2009). Highly connected networks often support redundant pathways of interaction, meaning the networks are more resilient to disturbance but often less specialized (Dunne *et al*., 2002; Liu *et al*., 2025). In contrast, modularity quantifies the extent to which species form clusters or “modules” of tightly associated taxa that interact more strongly within than between modules (Newman, 2006). Modular networks often emerge when specific microhabitats, substrates, or host associations constrain interactions, enhancing local stability but limiting cross-module links. Each of these network properties can offer valuable insight into the assembly of fungal-beetle networks and their potential responses to environmental change, yet our understanding of these dynamics remains limited.

In deadwood, properties of fungal and beetle networks differ among tree species due to variations in the abiotic and biotic context (Bouget *et al*., 2013; Wende *et al*., 2017; Kriegel *et al*., 2023). Species richness of macrofungi, lichens and beetles in Sweden are higher in conifers than in broadleaved trees except for oak (Sundberg *et al*., 2019). However, for the broadleaved trees like birch, have been shown to support a higher per-log species richness of wood-inhabiting fungi (and their associated saproxylic beetles) compared to coniferous deadwood (Abrego *et al*., 2016; Purhonen *et al*., 2020a). However, patterns of fungal species richness inferred from molecular identification varies. Some metabarcoding studies have reported equal or higher fungal richness in coniferous deadwood at the log scale (e.g. Purahong *et al*., 2018a), indicating that richness estimates depend on the context and methodological approach. Fungal and beetle community compositions differ among tree species and increasingly so with larger phylogenetic distance <u>(e.g. Purahong *et al*., 2018; Stokland et al 2012; Sundberg at al 2019)</u>. While tree species are therefore expected to differ in fungal–beetle interaction structure due to differences in tree species traits and associated species pools, empirical evidence directly linking tree species identity to differences in fungal–beetle network structure (e.g. modularity, nestedness) is still to be explored.

In addition to tree species, the deadwood type (e.g., left after forest enrichment or created as a restoration measure) can also influence the fungal-beetle network structure (Hjältén *et al*., 2012; Hägglund & Hjältén, 2018). For example, fire-created deadwood selects pyrophilous beetles and fungal assemblages, with members colonizing recently burned wood, but otherwise being rare in the landscape. This leads to highly specialized and modular networks. Felled trees (e.g., cut trees lying on the ground), on the other hand, retain moisture better than standing dead trees. They typically support diverse saproxylic fungal communities, which in turn attract many different beetle species (Purhonen *et al*., 2020b). Studies show that decomposition proceeds faster in felled deadwood and host a larger number of species than standing dead stumps of the same age (Lindhe *et al*., 2004), often resulting in higher network nestedness and connectance as many generalist beetles interact with multiple fungal species (Koivula & Vanha-Majamaa, 2020). Standing deadwood, such as high stumps and girdled trees, typically represent relatively dry deadwood habitats, which can limit early fungal colonization and, in turn, beetle establishment. When deadwood types are compared at a similar age, standing substrates therefore tend to host fewer fungal taxa than felled logs, especially during early to intermediate decomposition (Persiani *et al*., 2015; Sefidi & Etemad, 2015; Larsson Ekström *et al*., 2024a). However, these differences are not fixed over time: standing deadwood may later fall, after which moisture conditions increase and colonization by additional fungi and beetles can occur. Overall, changes in forest management practices that alter the relative availability of deadwood types, e.g., enrichment practices, are likely to shift the colonization patterns of specialist- and generalist fungal and beetle species and thereby influence the specialization, modularity, and connectivity of fungal–beetle networks.

To better understand how deadwood of different tree species and deadwood enrichment influence multitrophic interactions, we assessed the co-occurrence patterns between fungi and saproxylic beetles across deadwood tree species and deadwood types in a large-scale restoration experiment. Established in 2011, the experiment included 12 forest stands subjected to either *prescribed burning* (6 stands) or *artificial gap creation* (6 stands). Within each gap stand, four deadwood types, namely *felled logs*, *girdled*, *high stumps*, and *uprooted trees*, of Norway spruce (*Picea abies*), Scots pine (*Pinus sylvestris*) and downy birch (*Betula pubenscens*) were created to mimic natural disturbance regimes. Based on the extensive existing literature on deadwood fungal and beetle communities (e.g., Stokland *et al*., 2012), we expect that:

1. fungal and beetle community composition differ among tree species and deadwood types;
2. fungal and beetle species show structured (i.e., non-random) associations, with stronger co-occurrence patterns driven by tree species identity than by deadwood type;
3. tree species and, to lesser extent, deadwood type impact network properties such as nestedness, modularity, and specialization.

Specifically, for fungal–beetle network structure, we predict that:

3a) Tree species exert strong effects on network structure and fungal and beetle communities co-occurrence patterns, because differences in host traits lead to differences in species pools and thus community composition.
3b) Burned dead trees host distinct communities of fungi and beetles selected to fire-affected substrates, as fire and consequently burnt wood have ecologically and evolutionary acted as selective forces. We predicted high modularity and specialization, low nestedness, and relatively high co-occurrence patterns reflecting coupled community turnover between fungi and beetles in burned trees.
3c) Felled and uprooted deadwood support the highest species richness and the beetle fungal network with many generalist species due to more moist and favorable conditions for many fungi and fungal decomposition, hence also for many beetles. Consequently, we expected high connectance and nestedness due to high co-occurrence of generalist taxa.
3d) For girdled and high stump deadwood types, we expected that the drier microclimatic conditions will constrain fungal and beetle colonization, resulting in species-poor assemblages. Consequently, we anticipated high modularity, but low connectance.

## Materials and methods

### Study area

The study was conducted in the middle and northern boreal zones in northern Sweden (Ahti *et al.*, 1968; Fig. 1a), across 12 forest stands (six selected for prescribed burning and six for gap-cutting) selected for similarity in age, tree species composition, ground vegetation, and productivity. Stand information was provided by the landowner, with final selection based on field visits, but also collected during the following beetle sampling events (Fig.1a; Table S1). Stands (3.5–21 ha) were dominated by Scots pine (*Pinus sylvestris*) and/or Norway spruce (*Picea abies*), with scattered deciduous species including birch (*Betula pubescens*), aspen (*Populus tremula*), and goat willow (*Salix caprea*). Most sites were mesic dwarf-shrub types with *Vaccinium myrtillus* dominating the field layer, though some had moist or dry patches. To reduce variation, stand characteristics were standardized across burned and gap cuts: 80–160 years in age, 30–70% pine, 30–60% spruce, 5–20% deciduous, and 150–270 m³ ha⁻¹ in volume. All stands had a history of selective logging but were never clear-felled, resulting in reduced structural complexity compared to unmanaged forests (Stenbacka *et al*., 2010). However, they retain forest continuity and are suitable for ecological restoration. Due to effective fire suppression, natural fires are rare, highlighting the need for fire-based restoration. All stands are part of FSC-certified voluntary set-asides.

**Figure 1.**
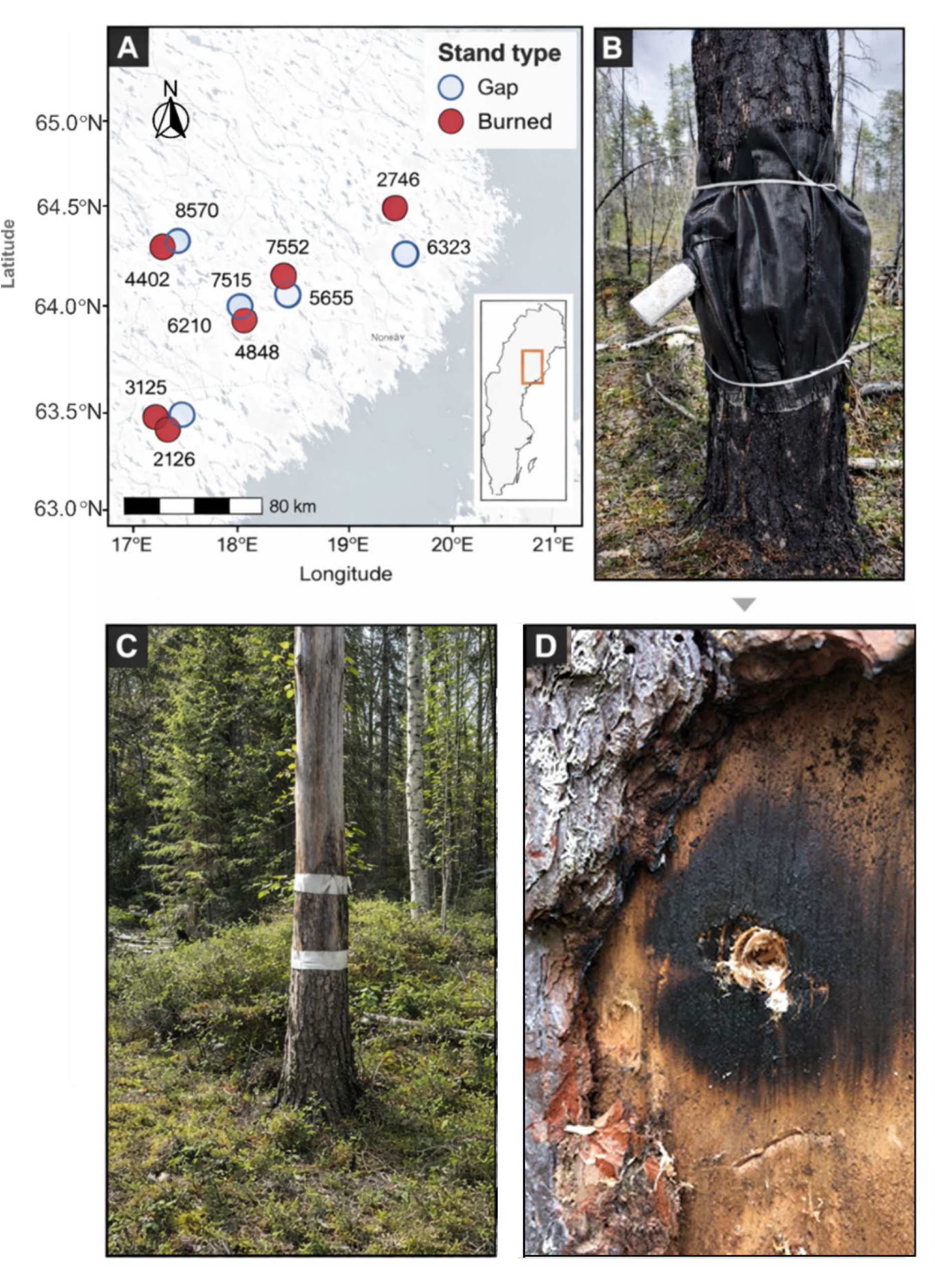
Panel (A) Location of the 12 forest stands in northern Sweden within the middle and northern boreal zones. Six stands were assigned to prescribed restoration burning and six to artificial gap creation with coarse woody debris enrichment. Final replication numbers for each deadwood type × tree species combination are provided in Table S2. Panel (B) Emergence traps used for beetle collection. Black polypropylene mesh was wrapped tightly around the trunk and sealed with foam and plastic straps. A collecting bottle containing propylene glycol (∼60%) was attached to capture emerging insects. Traps were installed in late May 2020 and collected in September 2020. Panel (C) shows a high stump after trap removal. Sampling was conducted at the same position as the insect trap placement. Panel (D) shows close-up of fungal sampling procedure from a burned deadwood tree with the drilling into the substrate to collect sawdust for DNA extraction. Bark was removed prior to drilling with knife, and ca. 20 cm^2^ of the wood surface sterilized using a gas burner. Four subsamples per deadwood were pooled into one composite sample.

### Experimental setup

In 2011, a deadwood restoration experiment was established across 12 forest stands, with six stands assigned to restoration burning and six to artificial gap creation with coarse woody debris (CWD) enrichment (Fig. 1a). Within each gap-cut stand, six 20-meter diameter canopy gaps per hectare were created using a standard harvester. In the gaps, four types of deadwood treatments were implemented to mimic natural disturbance: (i) felled logs (trees cut at the base and left as logs), (ii) high stumps (trees cut ∼3 m above ground), (iii) simulated windthrows (trees pushed over), and (iv) girdled trees (trees girdled at ∼3 m height and left standing; (Fig. 1a). All types of deadwood were created of pine and spruce trees. In addition, birch *(B. pubescens*) trees were felled down. Due to shortage of birches in the stands, no other types of birch deadwood could be created. In the burned stands we selected standing birch, pine and spruce trees that had died during or right after prescribed fire (Fig. 1a). In each study site, we aimed in including 5 replications of each deadwood type and tree species combinations. However, some replicates were lost due to sampling or processing issues, for example, failed beetle trap collections or unsuccessful fungal DNA amplifications. Thus, in total, final dataset consisted of 348 beetle community samples and 356 fungal community samples (see Table S2 for details on the final sample counts).

### Beetle collection and morphological identification

The beetles were collected using emergence traps that catch the insects emerging from the deadwood substrates and therefore give a representative sample of beetles inhabiting the substrate. The emergence traps consist of black polypropylene weed barrier wrapped around the trunk of the tree (Fig 1b). Traps were attached to the tree tightly with plastic straps, leaving a space of 30-40 cm between the straps. A foam rubber was used under the straps to seal the ends of the traps, and metal wires were used to keep the mesh apart from the tree trunk to allow air circulation under the mesh. A semi-transparent plastic bottle was attached to the side of the trap to collect the emerging insects. Emerging insects are drawn by light and will crawl into the collecting bottle. The preservative liquid was propylene glycol diluted to ca 60% and with a small amount of detergent to reduce the surface tension. The traps were set up in late May 2019 and collected in September 2019. The insects were sorted and identified to species level by a taxonomic expert, Dr. Petri Martikainen.

### Fungi sampling and DNA sequencing

During autumn 2019, fungi were sampled by drilling each deadwood substrate to collect sawdust for DNA extraction (Fig. 1cd). The samples were collected from each deadwood type from the same position where the insect emergence trap was placed. Before drilling, the bark layer was removed from a drilling point with knife, and ca. 20 cm^2^ of the wood surface sterilized using a gas burner. Four sawdust samples were collected from each unit and pooled into one collective sample. The samples were taken using a 10 mm diameter drill (Pasanen *et al*., 2018), which was sterilized with a blowtorch between samples.

The sawdust samples used for DNA extraction were freeze-dried in their collected 50 mL. DNA was extracted from 0.5 g sawdust using the NucleoSpin Soil kit (Macherey-Nagel, Düren, Germany) with SL1 buffer and enhancer SX. The fungal ITS2 region was amplified using the tagged primers gITS7 and a 3:1 mix of ITS4 and ITS4arch (Ihrmark *et al*., 2012; Clemmensen *et al*., 2016). PCRs were run in 50-µL reactions with 5 µL DNA template and 20–35 cycles, depending on amplification strength. Duplicate PCR products were pooled, purified, and sequenced at SciLifeLab/NGI (Uppsala, Sweden) on a PacBio RSII platform (Castaño *et al*., 2020). Sequences were quality filtered and clustered into fungal species hypotheses in SCATA using single-linkage clustering at 98.5% similarity, and fungal taxa were assigned against the UNITE database after excluding non-fungal sequences (Abarenkov *et al*., 2024). Full methodological details are given in Supporting Information Methods S1.

### Statistical analysis

All analyses were conducted in R 4.4.3 (R Core Team, 2025) with the packages *bipartite* (Dormann *et al*., 2025), *igraph* (Csárdi *et al*., 2026), *ggplot2* (Wickham, 2009), *vegan* (Oksanen *et al*., 2022)*, tidyverse* (Wickham & RStudio, 2023) and *patchwork* (Pedersen, 2025).

#### 1. Community composition and indicator species analyses

To explore the differences in fungal and beetle community composition among tree species and tree species and deadwood types, we performed PERMANOVA using function *adonis2* in the *vegan* package. First, we ran multivariate models, to explore the differences in community composition among three tree species. Second, we ran the model to explore the differences in community composition between felled and burned trees of birch trees. We also explored differences in fungal and beetle community composition among burned standing, girdled, high-stump, uprooted, and felled pine and spruce deadwood by conducting pairwise permutation-based MANOVAs (PERMANOVA) on Bray–Curtis dissimilarities using the function. *pairwise.perm manova* implemented in the *RVAideMemoire* package with 999 permutations and FDR correction for multiple testing. As the final step, we performed indicator species analysis using function *multipatt* in the package *indicspecies* to explore which species are associated with which tree species and tree species × deadwood types.

#### 2. Multivariate correlations

To assess the degree of multivariate correlations between fungal and beetle communities (e.g., whether fungal and beetle communities change in parallel across tree species and tree species and deadwood types), we calculated the RV coefficient (Escoufier, 1973) using the *RV.rtest* function in the *ade4* package (Dray *et al*., 2025). The RV coefficient is a multivariate generalization of the squared Pearson correlation, quantifying the coupling between two sets of community data. Values range from 0 (no correspondence) to 1 (complete concordance). Prior to analysis, both beetle and fungal abundance matrices were Hellinger-transformed (*decostand* in the *vegan* package) to standardize abundances and reduce the influence of highly dominant taxa. RV coefficients were computed separately for each tree species and tree species × deadwood type combination. The significance of RV values was tested with 999 permutations, where sample rows were randomly permuted in one dataset relative to the other. In practical terms, a high RV coefficient indicates that differences in fungal communities among tree species or deadwood types are associated with similar differences in beetle communities, whereas a low RV indicates weak or no links between the two groups.

#### 3. Network analysis

##### a) Network inference and construction

For the construction of the networks, co-occurrence matrices were constructed from the joint presence of taxa, providing a statistical approximation of potential associations. While such patterns can arise from shared ecological preferences (e.g., similar microhabitats, substrates, or decay stages), they may also reflect direct ecological interactions, particularly for fungivore beetles that rely on fungal tissues for feeding or oviposition. Networks were constructed using a full dataset, and all inferred co-occurrences were used to characterize overall network structure and to test whether their structure significantly differs from chance and to explore species-level roles.

Prior to network construction, dataset was filtered to retain: 1) OTUs and beetle species that were observed in at least three samples and with a total abundance greater than two; 2) OTUs (for fungi) and individuals for beetles identified to species level. We calculated three indices to quantitatively characterize the structure of the interaction networks: connectance (i.e. the proportion of possible links in the network that are realized), nestedness (a measure of the degree to which species interactions in a network are hierarchically organized, such that species with fewer interactions form subsets of the interaction partners of species with more interactions) and modularity. Connectance and nestedness were calculated using the functions *networklevel()* and *nestednodf()* from the *bipartite* package (Dormann *et al*., 2025), while modularity was calculated using *cluster_fast_greedy()* from the *igraph* package (Csárdi *et al*., 2026). Networks were constructed and analyzed separately for each tree species (pine, spruce, birch) and each tree species × deadwood type (e.g., burned standing, felled, girdled, high stump, uprooted) combination to capture type specific patterns. To explore whether all network indices significantly differ from random, we used two null model types to compare observed indices with simulated values (1000 times). We used r00 (*bipartite* package) and model2 null model types from the *vegan* package (Oksanen *et al*., 2022). The r00 model in *bipartite* package is a constrained randomization model. It creates randomized matrices while maintaining the degree distribution (row and column sums) of the observed network. *Model2* from the vegan package (Oksanen *et al*., 2022) randomizes the matrix based on species’ marginal totals and thus preserves row and column sums. For each null model, we generated 1000 randomized networks and calculated standardized effect sizes (z-scores) and probabilities for each index, providing a test of whether connectance, nestedness, and modularity significantly departed from chance (Table 2).

##### b) Species roles within networks

To assess the roles of individual taxa within the networks, we constructed fungus–beetle bipartite matrices separately for each tree species and deadwood type and explored species roles within those networks. In brief, for each network, we calculated species-level metrics using the function *specieslevel()* in the *bipartite* package (Dormann *et al*., 2025). To evaluate species’ structural roles, we identified network modules with *computeModules()* (*bipartite* package) and calculated within-module degree (z) and participation coefficient (c) using *czvalues*() following the thresholds proposed by (Guimerà & Nunes Amaral, 2005). The within-module degree reflects how strongly a species is connected to others within its own module, whereas the participation coefficient captures how evenly its interactions are distributed across different modules. Species with low z and low c were categorized as peripherals (few species links confined within their own module), those with low z but high c were assigned to connectors (e.g., those that are linking different modules together), those with high z but low c as module hubs (highly connected within a single module), and those with both high z and high c as network hubs (e.g., species which are generalists, e.g. potentially interact with species within the module and across the modules).

## Results

### 1. Fungal and beetle community composition and indicator species differ among tree species and deadwood types

For fungi, after filtering and quality control, 2,078 fungal OTUs were described from three tree species and five deadwood types. Of the 2,078 fungal OTUs, 846 (40.7%), 740 (35.6%), 656 (31.6%), 578 (27.8%), 538 (25.9%), and 429 (20.6%) could be assigned to phylum, class, order, family, genus, and species levels, respectively, when excluding OTUs labelled “Unknown/Unidentified”. As for beetles, in total, 216 beetle species were identified. Birch had 72 species on burned standing and 60 on felled trees. In pine and spruce, respectively, 59 and 50 beetle species were found in burned standing trees, 44 and 59 in felled, 31 and 35 in girdled, 63 and 45 in high stumps, and 49 and 48 in uprooted trees.

#### 1.1. Tree species and to lesser extent deadwood type structure fungal and beetle community composition

Fungal community composition differed significantly among tree species, with tree species explaining 10.4% of the variation (Fig. 2a and Table 1). Within birch, fungal community composition differed between burned standing and felled deadwood (Fig. 2b and Table 1). For pine and spruce, fungal community composition varied significantly with tree species, deadwood type and their interaction (Fig.2c and Table 1). Pairwise comparisons showed that uprooted and felled deadwood differed from high stumps, girdled trees, and burned standing trees (*P_fdr_* ≤ 0.001; Fig. 2c). However, felled and uprooted deadwood had similar communities (for spruce *P_fdr_* = 0.110 and pine *P_fdr_* = 0.529, respectively; Fig. 2c). In pine, fungal community composition on burned standing and girdled trees differed marginally (Fig. 2c).

**Figure 2.**
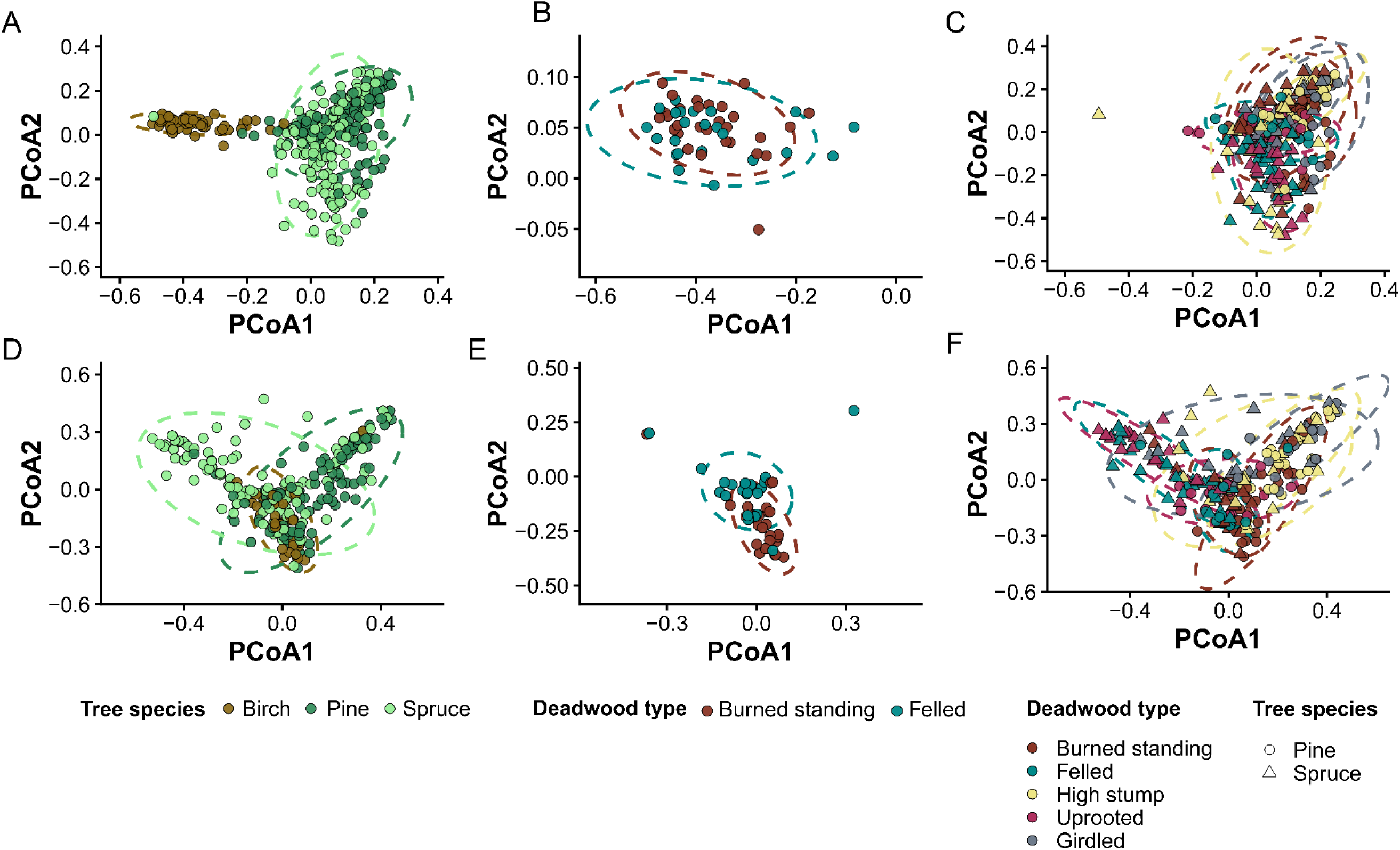
Principal coordinate analysis (PCoA) of Bray-Curtis dissimilarities showing the differences in fungal (A-C) and beetle community composition (D-F) among tree species and tree species × deadwood type combination. Each point represents one fungal or beetle sample, and ellipses represent 95% confidence regions based on the multivariate normal approximation of the point distribution. Each ellipse colour represents either a tree species in panels A and D or a deadwood type in panels B, C, E and F.

**Table 1.** Variation in fungal and beetle composition across deadwood tree species and deadwood types. Statistical analysis of fungal and beetle communities revealed significant differences in composition across tree species (birch, pine and spruce) and deadwood types (), as determined by permutational multivariate analysis of variance (PERMANOVA) on Bray-Curtis dissimilarities. F: pseudo-F value, df: degrees of freedoms, R^2^: r-squared, *P*: *p*-value.

|  | **Models** | **Factor** | **Factor levels** | **df** | **F** | **R<sup>2</sup>** | ***P*** |
| --- | --- | --- | --- | --- | --- | --- | --- |
| **Fungi** | 1) All samples | Tree species | Birch, spruce and pine | 2 | 17.08 | 0.104 | 0.0001 |
|  | 2) Birch | Deadwood type | Felled and burned standing | 1 | 2.57 | 0.049 | 0.001 |
|  | 3) Pine and Spruce | Tree species | Pine and spruce | 1 | 9.96 | 0.036 | 0.001 |
|  |  | Deadwood type | Felled, uprooted, high stump, girdled & burned standing | 4 | 4.58 | 0.067 | 0.001 |
|  |  | Tree species × deadwood type |  | 4 | 2.79 | 0.041 | 0.001 |
| **Beetles** | 1) All samples | Tree species | Birch, spruce and pine | 2 | 6.81 | 0.045 | 0.0001 |
|  | 2) Birch | Deadwood type | Felled and burned standing | 1 | 3.43 | 0.064 | 0.001 |
|  | 3) Pine and Spruce | Tree species | Pine and spruce | 1 | 6.10 | 0.092 | 0.001 |
|  |  | Deadwood type | Felled, uprooted, high stump, girdled & burned standing | 4 | 7.44 | 0.028 | 0.001 |
|  |  | Tree species × deadwood type |  | 4 | 4.44 | 0.147 | 0.001 |

Beetle community composition also differed among tree species (Fig. 2d and Table 1). Within birch, communities differed between burned standing and felled deadwood (Fig. 2e and Table 1). For pine and spruce, beetle communities varied with tree species, deadwood type and their interaction (Fig. 2f and Table 1). Furthermore, the pairwise comparisons among spruce deadwood types showed beetle community composition differed significantly among most deadwood treatments (*P_fdr_* ≤ 0.001; Fig. 2f). Significant differences were detected between high-stump and felled deadwood, as well as between felled and both girdled and burned standing deadwood (Fig. 2f). In contrast, beetle community composition on felled and uprooted spruce trees were similar (*P_fdr_* = 0.891; Fig. 2f). For pine, beetle community composition differed significantly among most deadwood types, except for felled and uprooted (*P_fdr_* = 0.502) and girdled and high stumps (*P_fdr_* = 0.725; Fig. 2f).

#### 1.2. Fungal and beetle species-level deadwood type specialization within tree species and strong differentiation between standing (burned, high stump and girdled) and fallen (felled and uprooted) deadwood types

Indicator species analyses revealed clear differentiation among tree species and weaker differentiation among deadwood types (Fig. 3). The species-level patterns matched genus-level shifts in species frequencies (Fig. S1).

**Figure 3.**
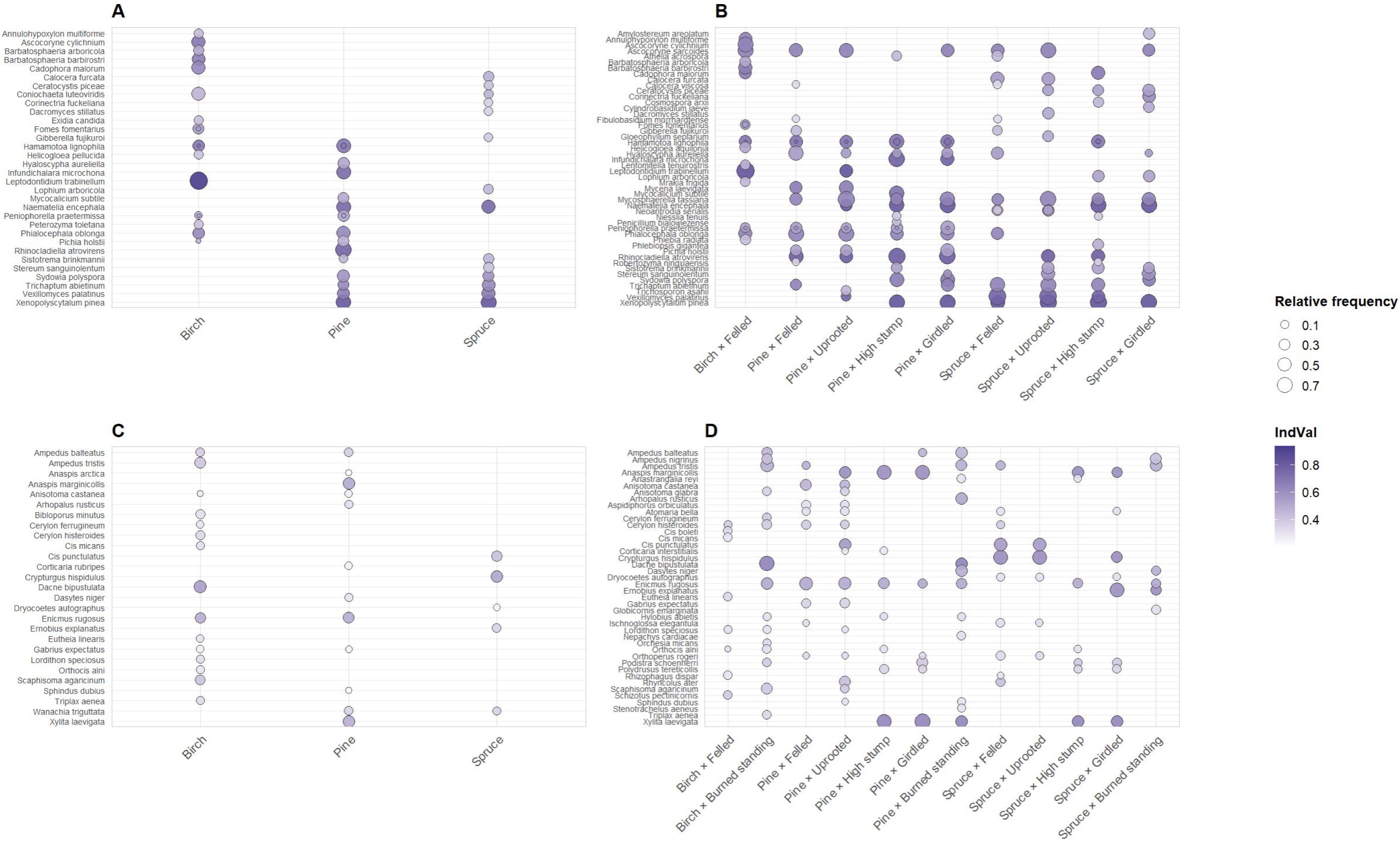
Indicator taxa across tree species and restoration treatments. Bubble plots show taxa identified as significant indicators for (A–B) fungi and (C–D) beetles. Panels on the left summarize indicators by tree species (birch, pine, spruce), whereas panels on the right show indicators for each tree species × deadwood type combination. Bubble color represents indicator value (IndVal; darker violet = stronger association), and bubble size represents the taxon’s relative frequency (i.e., proportion of samples in which the taxon was detected) within each group. For each group, taxa ranked by IndVal and ordered alphabetically on the y-axis.

For fungi, indicator species differed among tree species (Fig. 3a). Birch was characterized by the ascomycetes *Ascocoryne cylichnium*, *Cadophora malorum*, and *Leptodontidium trabinellum*, together with unidentified taxa in *Bartabtosphaeria* and *Coniochaeta*. Pine and spruce were associated with conifer-deadwood fungi, including *Xenopolyscytalum pinea*, *Vexillomyces palatinus*, *Rhinocladiella atrovirens*, and *Trichaptum abietinum*; *Corinectria fuckeliana*, *Gibberella fujikuroi*, and *Ceratocystis piceae* were also significant spruce indicators. These patterns matched genus-level composition (Fig. S2c): birch was dominated by *Cadophora* and *Leptodontidium*, whereas conifer deadwood had higher contributions of *Trichaptum*, *Vexillomyces*, *Naematelia*, and *Xenopolyscytalum*. Deadwood types yielded few indicator species (Fig. 3b); felled spruce was characterized by *Stereum sanguinolentum*, *Sydowia polyspora*, *Naematelia encephala*, and *Vexillomyces palatinus*.

Beetle indicator species (Fig. 3cd) also showed significant tree species associations. Birch was most strongly associated with *Dacne bipustulata* and *Enicmus rugosus*, while pine indicators included taxa such as *Xylita laevigata* and *Anaspis marginicollis*. Spruce was characterized by indicators consistent with conifer deadwood and bark-associated species, including fungivore *Cis punctulatus* and cambivore *Crypturgus hispidulus* (Fig. 3c). Several beetle species showed clear substrate-specificity within tree species (Fig. 3d). Several elaterids and fungivorous beetle species, including *Ampedus* spp. (e.g., *A. tristis*, *A. nigrinus*, *A. balteatus*) and *Dacne bipustulata* were significant indicator species of burned-standing and felled birchs, alongside fungivorous *Enicmus rugosus* and *Scaphisoma agaricinum* (Fig. 3d). Pine substrates showed strong differentiation among deadwood types; *Anaspis marginicollis* and *Xylita laevigata* (Fig. 3d) were indicators of girdled pines and high stumps, whereas *Arhopalus rusticus* and *Cerylon histeroides* were indicators of felled pine, *Dasytes niger* of burned standing and *Rhyncolus ater* of uprooted pine. For spruce, *Cis punctulatus* and *Crypturgus hispidulus* were indicator species of burned standing, felled and uprooted spruces.

Finally, when exploring the indicator taxa across deadwood types, fungi like *Phialocephala oblonga*, *Phlebopsis gigantea*, *Stereum sanguinolentum* and *Vexillomyces palatinus* were indicator species of high stump, while *Naematelia encephala* and *Nieslia sp*. of girdled trees (Fig. S2). As for beetles, *Ampedus balteatus*, *Ampedus nigrinus* and *Ampedus tristis* were indicator species of burned standing wood, while *Xylita laevigata* was an indicator species of high stumps and girdled trees (Fig. S2). In addition, a few beetle species were indicators of more than one deadwood type. For example, *Dacne bipustulata* was an indicator of burned standing deadwood, but also uprooted trees, while *Dasytes niger* was an indicator species of burned standing and girdled trees (Fig. S2).

### 2. Fungal–beetle community links are substrate-specific and strongest in burned and girdled conifer deadwood

The strength of multivariate covariation between fungal and beetle community composition, quantified using the RV coefficient, varied among tree species and deadwood types (Fig. 4). At the tree species level, no significant multivariate correlations between fungi and beetles were observed for birch (RV = 0.47, P = 0.388, n = 47; Fig. 4a), while significant multivariate correlations between beetle and fungal communities were detected for pine (RV = 0.39, P = 0.001; Fig. 4b) and spruce (RV = 0.34, P = 0.001, n = 125; Fig. 4c).

**Figure 4.**
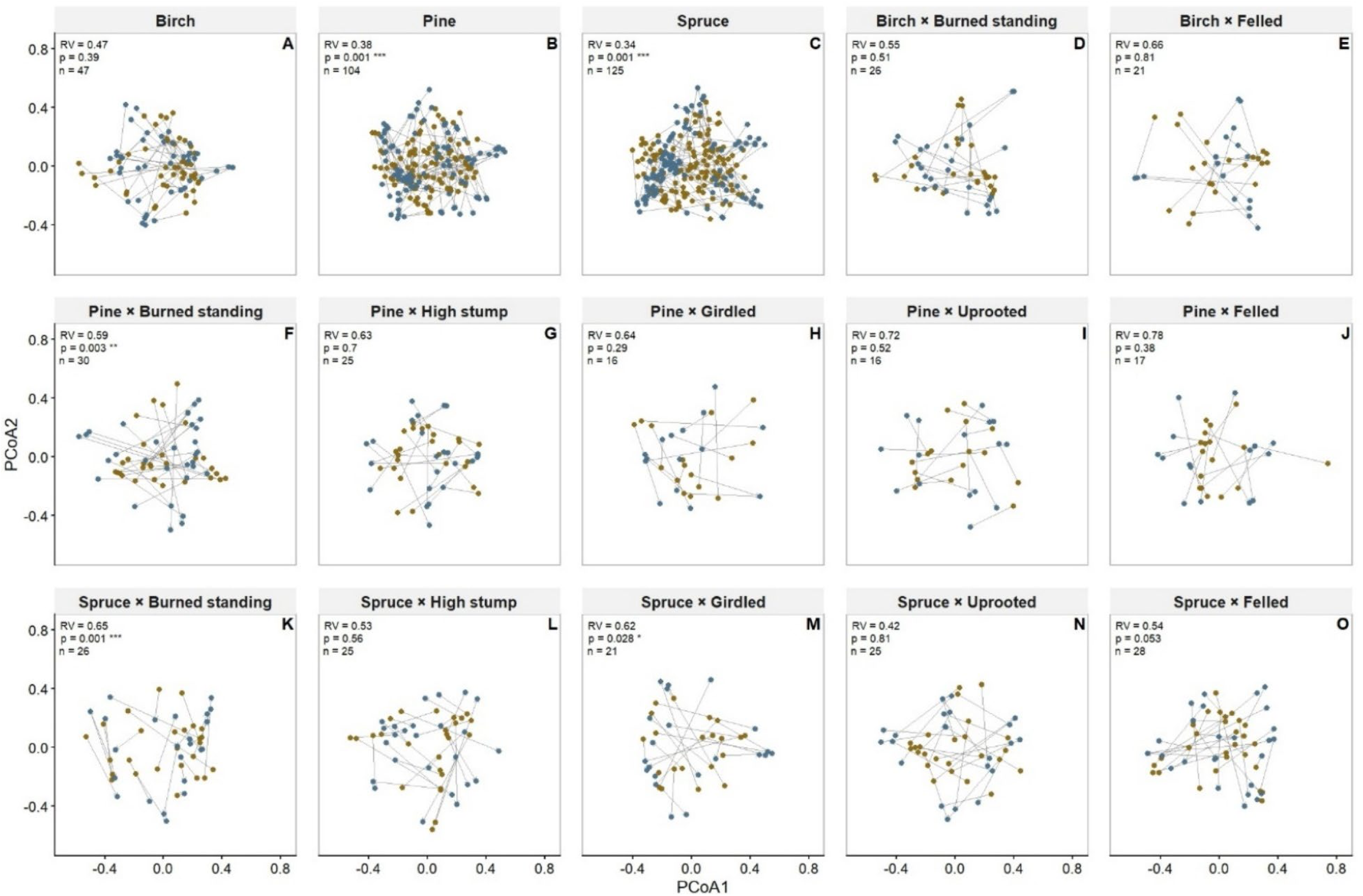
Multivariate correlations between fungal and beetle community composition across tree species (Panels A-C) and deadwood types (Panels D-O). Each panel represents one combination of tree species (rows: birch, pine and spruce) and deadwood type (columns: burned standing, felled, girdled, high stump, uprooted). Points correspond to ordination scores from Principal Coordinates Analysis (PCoA) based on Bray–Curtis dissimilarities of Hellinger-transformed community matrices. Arrows connect the fungal sample (olive green) of each tree to its corresponding beetle (dark blue) community, illustrating the degree of associations between the two groups within the same sample. Panels are annotated with the RV coefficient (a measure of multivariate correlation between species matrices) and the associated p-value from a permutation-based Procrustes test (`protest`). RV close to 0 indicates little or no multivariate correlations between species communities, while values closer to 1 indicate strong multivariate correlations. Significant associations are indicated with asterisks (*p* < 0.05 *, *p* < 0.01 **). Short, parallel arrows indicate strong agreement between fungal and beetle community structures (high multivariate correlation), while long, divergent arrows reflect weak correlation.

For the tree species × deadwood type interaction, the strength and significance of beetle–fungal multivariate correlations differed among deadwood types. For birch, no significant multivariate correlations were detected between beetle and fungal communities in either burned or felled trees (Fig. 4de). For pine, a significant correlation was detected for burned standing trees (RV = 0.59, P = 0.003, n = 30; Fig. 4f), whereas other pine deadwood types (girdled, high stump, felled, uprooted) were not statistically significant (Fig. 4g-j). As for the spruce, significant correlations were observed for burned standing trees (RV = 0.65, P = 0.001, n = 26; Fig. 4k) and girdled logs (RV = 0.62, P = 0.028, n = 21; Fig. 4m). Burned standing and girdled spruces showed strong multivariate correlations between fungal and beetle communities.

### 3. Fungal–beetle networks are modular and shaped by tree species and deadwood type Network level

Across all tree species and deadwood types, beetle–fungal co-occurrence networks showed low nestedness and high modularity compared to the null expectations (Table 2). Observed connectance closely matched the fixed–fixed (r00) null and only slightly exceeded the probabilistic null, suggesting that link density largely reflects species’ marginal totals, meaning that the number of cooccurrences is driven by how frequently beetle and fungal species occur overall, rather than by strong non-random associations between partners. All networks were less nested than expected, with strongly negative z-scores (Table 2). In contrast, networks were more modular and less nested than the null model with randomized interactions, with large positive z-scores, especially for pine and spruce. The network of fungi and beetles had the highest degree of modularity on felled birches, felled spruces, uprooted pine, pine and spruce high stumps (Table 2). Instead, the lowest modularity was detected for girdled spruce, burned spruce and burned birch (Table 2).

**Table 2.** Summary of network metrics for beetle–fungal co-occurrence networks across tree species and deadwood types. . For each network (defined by tree species and tree species × type combination), we report the observed mean values of three structural metrics, such as connectance, nestedness (NODF) and modularity, alongside the corresponding means and standard deviations of two null model simulations: (1) the fixed–fixed (r00) model and (2) a probabilistic model based on row and column marginal totals. Z-scores and p-values indicate the statistical deviation of the observed networks from null expectations, with significance levels marked as follows: *p ≤ 0.001, p ≤ 0.01, p ≤ 0.05, and ns = not significant.

| Network type | Connectance |  |  |  |  | Nestedness<br>(NODF) |  |  |  | Modularity |  |  |  |
| --- | --- | --- | --- | --- | --- | --- | --- | --- | --- | --- | --- | --- | --- |
|  | Null<br>model | Obs.<br>value | Mean<br>(sd) | z-score | P-value | Obs.<br>value | Mean<br>(sd) | z-score | P-value | Obs.<br>value | Mean<br>(sd) | z-score | P-value |
| 1. Birch | nm1 | 0.377 | 0.377<br>(0) | - | 1 | 33.12 | 82.2<br>(1.19) | -41.22 | < **0.001** | 0.168 | 0.123<br>(0.002) | 24.82 | < **0.001** |
|  | nm2 | 0.377 | 0.377<br>(0) | 0.003 | 1 | 33.12 | 61.36<br>(1.28) | -22.01 | < **0.001** | 0.168 | 0.118<br>(0.003) | 17.33 | < **0.001** |
| 2. Pine | nm1 | 0.302 | 0.302<br>(0) | - | 1 | 28.94 | 85.14<br>(0.79) | -71.42 | < **0.001** | 0.180 | 0.108<br>(0.001) | 50.21 | < **0.001** |
|  | nm2 | 0.302 | 0.301<br>(0.003) | 0.044 | 1 | 28.94 | 61.05<br>(0.93) | -34.56 | < **0.001** | 0.180 | 0.103<br>(0.002) | 37.32 | < **0.001** |
| 3. Spruce | nm1 | 0.235 | 0.235<br>(0) | - | 1 | 28.58 | 79.18<br>(0.89) | -56.60 | < **0.001** | 0.199 | 0.128<br>(0.002) | 46.97 | < **0.001** |
|  | nm2 | 0.235 | 0.235<br>(0.002) | 0.183 | 1 | 28.58 | 57.14<br>(0.82) | -35.02 | < **0.001** | 0.199 | 0.123<br>(0.002) | 42.95 | < **0.001** |
| 4. Birch<br>(Burned<br>standing) | nm1 | 0.243 | 0.243<br>(0) | - | ns | 18.1 | 76.1<br>(1.60) | -36.2 | < **0.001** | 0.211 | 0.167<br>(0.002) | 20.7 | < **0.001** |
|  | nm2 | 0.243 | 0.242<br>(0.004) | 0.07 | ns | 18.1 | 51.5<br>(1.85) | -18.1 | < **0.001** | 0.211 | 0.159<br>(0.003) | 15.8 | < **0.001** |

| Network type |  | Connectance |  |  |  | Nestedness<br>(NODF) |  |  |  | Modularity |  |  |  |
| --- | --- | --- | --- | --- | --- | --- | --- | --- | --- | --- | --- | --- | --- |
| 5. Birch (Felled) | nm1 | 0.209 | 0.209 | - | ns | 16.6 | 73.8 | -43.3 | < 0.001 | 0.231 | 0.153 | 33.5 | < 0.001 |
|  |  |  | (0) |  |  |  | (1.32) |  |  |  | (0.002) |  |  |
|  | nm2 | 0.209 | 0.210 | -0.32 | ns | 16.6 | 49.4 | -22.3 | < 0.001 | 0.231 | 0.145 | 59.4 | < 0.001 |
|  |  |  | (0.003) |  |  |  | (1.47) |  |  |  | (0.001) |  |  |
| 6. Spruce (Burned standing) | nm1 | 0.204 | 0.204 | - | ns | 15.1 | 73.7 | -48.4 | < 0.001 | 0.192 | 0.142 | 23.8 | < 0.001 |
|  |  |  | (0) |  |  |  | (1.21) |  |  |  | (0.002) |  |  |
|  | nm2 | 0.204 | 0.205 | -0.27 | ns | 15.1 | 49.1 | -26.3 | < 0.001 | 0.192 | 0.142 | 23.8 | < 0.001 |
|  |  |  | (0.003) |  |  |  | (1.30) |  |  |  | (0.002) |  |  |
| 7. Spruce (Felled) | nm1 | 0.154 | 0.154 | - | ns | 13.1 | 64.9 | -80.3 | < 0.001 | 0.231 | 0.15 | 80.5 | < 0.001 |
|  |  |  | (0) |  |  |  | (0.65) |  |  |  | (0.001) |  |  |
|  | nm2 | 0.154 | 0.154 | - | ns | 13.1 | 41.5 | -41.7 | < 0.001 | 0.231 | 0.143 | 58.1 | < 0.001 |
|  |  |  | (0.001) |  |  |  | (0.68) |  |  |  | (0.002) |  |  |
| 8. Spruce (Uprooted) | nm1 | 0.154 | 0.154 | - | ns | 14.2 | 64.9 | -80.3 | < 0.001 | 0.231 | 0.15 | 80.5 | < 0.001 |
|  |  |  | (0) |  |  |  | (0.65) |  |  |  | (0.001) |  |  |
|  | nm2 | 0.154 | 0.154 | - | ns | 14.2 | 41.5 | -41.7 | < 0.001 | 0.231 | 0.143 | 58.1 | < 0.001 |
|  |  |  | (0.001) |  |  |  | (0.68) |  |  |  | (0.002) |  |  |
| 9. Spruce (High stump) | nm1 | 0.154 | 0 | - | ns | 13.1 | 64.6 | -103 | < 0.001 | 0.228 | 0.150 | 47.1 | < 0.001 |
|  |  |  |  |  |  |  | (0.50) |  |  |  | (0.002) |  |  |
|  | nm2 | 0.154 | 0 | - | ns | 13.1 | 41.8 | -39.1 | < 0.001 | 0.228 | 0.143 | 60.3 | < 0.001 |
|  |  |  |  |  |  |  | (0.74) |  |  |  | (0.001) |  |  |
| 10. Spruce (Girdled) | nm1 | 0.240 | 0.240 | - | ns | 15.1 | 74.0 | -44.4 | < 0.001 | 0.194 | 0.150 | 17.5 | < 0.001 |
|  |  |  | (0) |  |  |  | (1.33) |  |  |  | (0.002) |  |  |
|  | nm2 | 0.240 | 0.240 | 0.02 | ns | 15.1 | 48.2 | -28.1 | < 0.001 | 0.194 | 0.142 | 18.6 | < 0.001 |
|  |  |  | (0.003) |  |  |  | (1.18) |  |  |  | (0.003) |  |  |
| 11. Pine (Burned standing) | nm1 | 0.240 | 0.240 | - | ns | 20.0 | 79.6 | -57.0 | < 0.001 | 0.195 | 0.136 | 34.8 | < 0.001 |
|  |  |  | (0) |  |  |  | (1.05) |  |  |  | (0.002) |  |  |
|  | nm2 | 0.240 | 0.240 | 0.02 | ns | 20.0 | 54.2 | -33.0 | < 0.001 | 0.195 | 0.139 | 33.6 | < 0.001 |
|  |  |  | (0.003) |  |  |  | (1.03) |  |  |  | (0.002) |  |  |
| **12. Pine (Felled)** | *nm1* | 0.240 | 0.240 | - | ns | 20.0 | 79.5 | -52.1 | < **0.001** | 0.196 | 0.136 | 28.0 | < **0.001** |
|  |  |  | (0) |  |  |  | (1.14) |  |  |  | (0.002) |  |  |
|  | *nm2* | 0.240 | 0.240 | 0.05 | ns | 20.0 | 54.5 | -41.1 | < **0.001** | 0.196 | 0.129 | 31.8 | < **0.001** |
|  |  |  | (0.003) |  |  |  | (0.84) |  |  |  | (0.002) |  |  |
| **13. Pine (Uprooted)** | *nm1* | 0.20 | 0.20 | - | ns | 17.7 | 75.5 | -92.3 | < **0.001** | 0.22 | 0.13 | 77.0 | < **0.001** |
|  |  |  | (0) |  |  |  | (0.63) |  |  |  | (0.001) |  |  |
|  | *nm2* | 0.20 | 0.20 | -0.28 | ns | 17.7 | 49.5 | -32.8 | < **0.001** | 0.22 | 0.12 | 81.2 | < **0.001** |
|  |  |  | (0.002) |  |  |  | (0.97) |  |  |  | (0.001) |  |  |
| **14. Pine (High stump)** | *nm1* | 0.240 | 0.240 | 0.02 | ns | 20.0 | 79.3 | -52.5 | < **0.001** | 0.195 | 0.136 | 41.4 | < **0.001** |
|  |  |  | (0.003) |  |  |  | (1.13) |  |  |  | (0.001) |  |  |
|  | *nm2* | 0.240 | 0.240 | - | ns | 20.0 | 53.8 | -32.3 | < **0.001** | 0.195 | 0.129 | 47.4 | < **0.001** |
|  |  |  | (0) |  |  |  | (1.04) |  |  |  | (0.001) |  |  |
| **15. Pine (Girdled)** | *nm1* | 0.243 | 0.243 | - | ns | 19.4 | 81.4 | -73.8 | < **0.001** | 0.187 | 0.117 | 37.9 | < **0.001** |
|  |  |  | (0) |  |  |  | (0.84) |  |  |  | (0.002) |  |  |
|  | *nm2* | 0.243 | 0.243 | -0.308 | ns | 19.4 | 54.6 | -43.4 | < **0.001** | 0.187 | 0.112 | 39.4 | < **0.001** |
|  |  |  | (0.003) |  |  |  | (0.81) |  |  |  | (0.002) |  |  |

#### Species roles

Across tree species, fungal–beetle networks were dominated by peripheral taxa, with most species showing low within- and among-module connectivity and occurring in a single module (Fig. 5; Table S3). Connector species were rare but occurred in all three tree-species networks. In pine, these included the beetles *Xylita laevigata* and *Anaspis marginicollis*, and the fungi *Orbilia*, *Phlebiopsis gigantea*, and *Coniochaeta*; in spruce, connectors included the beetles *Cis punctulatus*, *Ampedus tristis*, and *Crypturgus hispidulus*, and the fungi *Neoantrodia serialis*, *Gloeophyllum sepiarium*, and *Calocera furcata* (Fig. 5; Table S3). Module and network hubs were rarer still and occurred only among fungi, including *Sydowia polyspora* and *Niesslia tenuis* as module hubs in spruce, and *Naematelia encephala* and *Serpula himantioides* as network hubs (Table S3).

**Figure 5.**
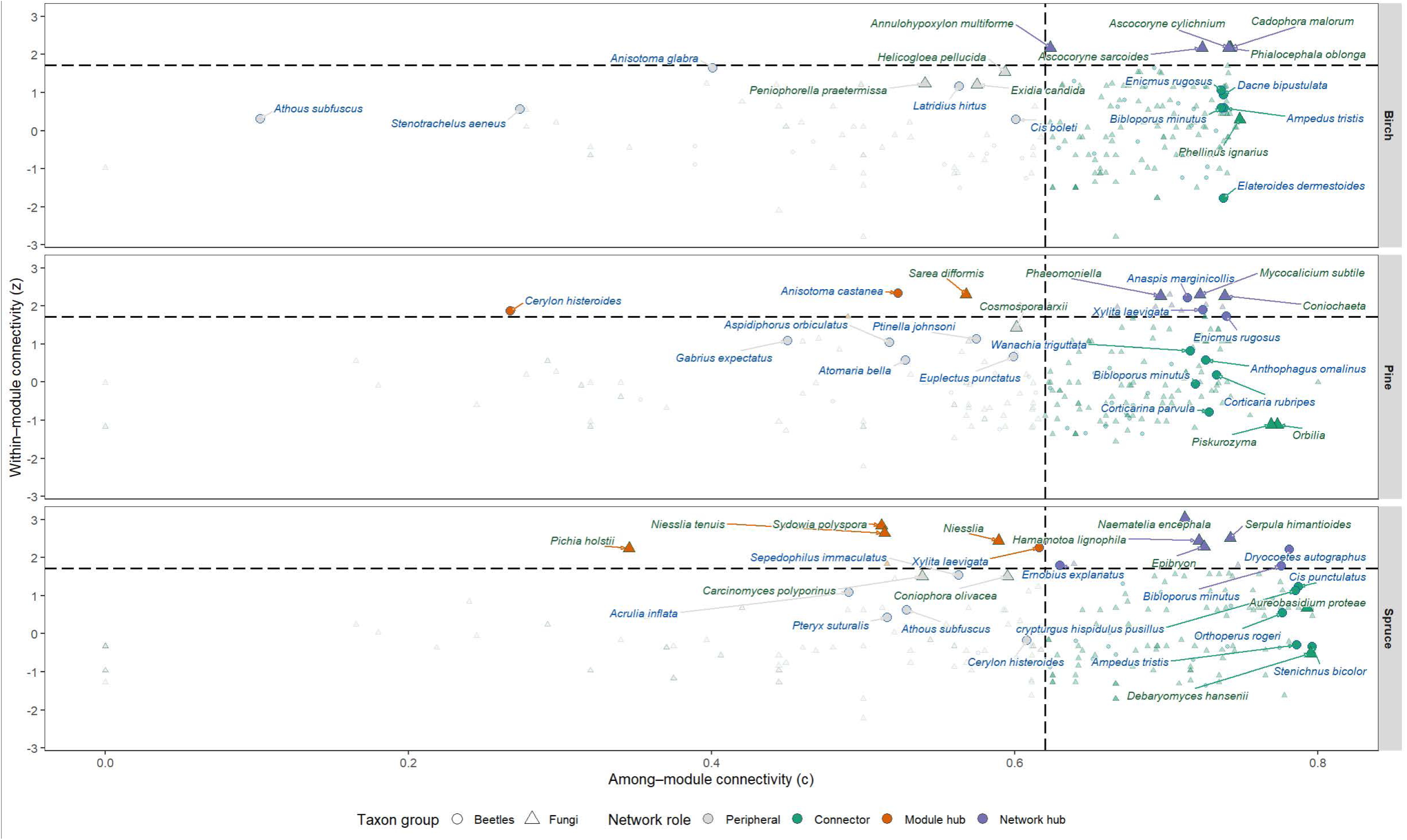
Network roles of beetle and fungal species across tree species. Scatterplots show the within-module connectivity (z) and among-module connectivity (c) for individual beetle (circles, dark blue color of the label) and fungal (triangles, olive green color of the label) species in bipartite co-occurrence networks constructed for each combination of tree species (birch, pine, spruce; rows) and deadwood treatment (burned standing, felled, girdled, high stump, uprooted; columns). Species are positioned according to their topological roles in the network: peripheral (grey fill; c ≤ 0.62, z ≤ 2.5), connector (gold fill; c > 0.62, z ≤ 2.5), and module hub (blue fill; z > 2.5). Species with low z and low c were categorized as peripherals (few species links confined within their own module), those with low z but high c as connectors (potentially linking multiple modules), those with high z but low c as module hubs (highly connected within a single module), and those with both high z and high c as network hubs (e.g., species which are generalists, e.g. potentially interact with species within the module and across the modules). Dashed vertical and horizontal lines indicate classification thresholds. Species labels indicate the top 5 nodes with the highest degree per panel.

Across tree species × deadwood type combinations, peripheral species dominated all networks, but the occurrence and identity of connectors and hubs varied among deadwood types within the same tree species (Fig. 6). Felled and uprooted deadwood supported more connectors and hub taxa across birch, pine, and spruce. In these deadwood types, fungal module hubs such as *Phaeomoniella* spp. and *Naematelia encephala* (a well-known parasite on *Stereum sanguiolentum;* Roberts, 1999; Pippola & Kytöviita, 2009) were frequent, especially in pine and spruce, forming within-module clusters often linked to multiple beetle species (Fig. 6; Table S3). Connector roles were also more common in lying deadwood, and included beetles such as *Gabrius expectatus* and *Cerylon histeroides*, and fungi such as *Cosmospora* spp. and *Sydowia polyspora*. In contrast, standing deadwood (high stumps and girdled trees) was dominated by peripheral taxa, particularly in pine and spruce, with most fungi and beetles restricted to single modules. Burned standing deadwood likewise contained few connectors or hubs despite supporting several specialized fungi and beetles (Fig. 6).

**Figure 6.**
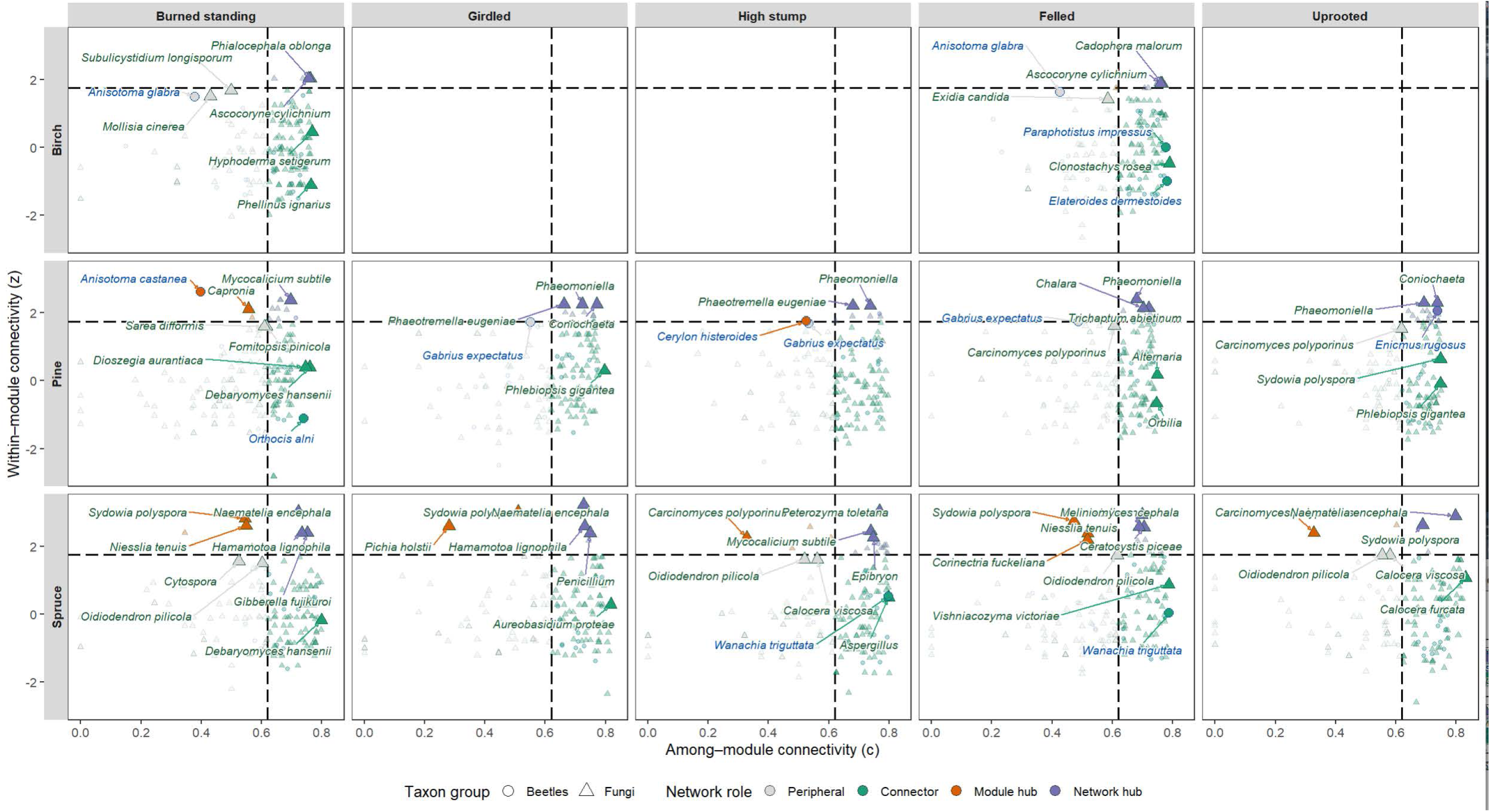
Network roles of beetle and fungal species across tree species and deadwood types. Scatterplots show the within-module connectivity (z) and among-module connectivity (c) for individual beetle (circles, dark blue color of the label) and fungal (triangles, olive green color of the label) species in bipartite co-occurrence networks constructed for each combination of tree species (birch, pine, spruce; rows) and deadwood treatment (burned standing, felled, girdled, high stump, uprooted; columns). Species are positioned according to their topological roles in the network: peripheral (grey fill; c ≤ 0.62, z ≤ 2.5), connector (gold fill; c > 0.62, z ≤ 2.5), and module hub (blue fill; z > 2.5). Species with low z and low c were categorized as peripherals (few species links confined within their own module), those with low z but high c as connectors (potentially linking multiple modules), those with high z but low c as module hubs (highly connected within a single module), and those with both high z and high c as network hubs (e.g., species which are generalists, e.g. potentially interact with species within the module and across the modules). Dashed vertical and horizontal lines indicate classification thresholds. Species labels indicate the top 5 nodes with the highest degree per panel.

## Discussion

Our results show that while tree species are associated with different fungal and beetle community composition and indicator taxa, some of those differences are also attributed to the deadwood enrichment types. The biggest differences in fungal and beetle species community composition were detected between standing (girdled and high stumps) and lying (felled and uprooted) deadwood types. Burned standing deadwood also differed in fungal and beetle communities from other standing deadwood types. The strength of multivariate covariation between fungal and beetle community composition varied among tree species and deadwood types: significant multivariate correlations were detected for pine and spruce, but not for birch. When tree species and deadwood type were considered jointly, significant correlations were restricted to burned standing pine, spruce, and girdled spruce. Importantly, all fungal–beetle networks were less nested and more modular than expected by chance, meaning that species were organized into small, distinct groups of co-occurring taxa rather than around a few generalists. This pattern was strongest in conifer deadwood, especially felled spruce and uprooted pine, where the networks were most strongly compartmentalized. Taken together, these results show that fungal–beetle co-occurrences in deadwood are highly structured, strongly dependent on deadwood species and types, but also partly structured by stochasticity. In practical terms, this means that managing a diversity of tree species and deadwood types is essential not only to maintain species community compositions, but also to preserve the specialized multitrophic networks.

### Fungal and beetle community composition and indicator species differ among tree species and to lesser extent among deadwood types

Consistent with our prediction, tree species, birch, spruce and pine, differed in fungal and beetle communities they harbored. This is no surprise given the number of studies in the past few decades that have shown the strong effect of tree species and, in particular, differences between broadleaf and coniferous trees (Gibb *et al*., 2005; Stokland *et al*., 2012; Hägglund & Hjältén, 2018; Sundberg *et al*., 2019; Löfroth *et al*., 2023). The strong effect of tree species in our study agrees particularly well with previous work on wood-inhabiting fungi. For example, Lepinay *et al*. (2022) found that fungal community composition in deadwood was driven primarily by tree species and wood chemistry, even across time, highlighting that host-tree identity remains a major filter throughout decomposition. Results on beetle community composition align with this pattern and likewise support our prediction. Many saproxylic beetles show strong host-tree associations, and previous studies have shown that tree species composition can be a major determinant of beetle abundance, richness and community composition (Stenbacka *et al*., 2010; Hjältén *et al*., 2012; Müller *et al*., 2015; Edelmann *et al*., 2022). Importantly, although tree species explained 10% of the variation in fungal and 6% in beetle community composition, a large proportion of variation remained unexplained. Such unexplained variation likely reflects differences in microclimatic conditions, bark cover, wood diameter, decay stage, local fungal and beetle priority effects, succession, stochasticity and connectivity among the deadwood trees all of which can vary among individual deadwood units even within the same tree species and deadwood type (Johansson *et al*., 2017; Dahlberg *et al*., 2025).

Deadwood type acted as a secondary but an important factor influencing the fungal and beetle community composition. One of the results to highlight is the difference between standing deadwood types (girdled, high stumps, and in many cases burned standing trees) and lying deadwood (felled and uprooted), with lying trees often supporting similar fungal and beetle assemblages, different from standing trees. This suggests that the physical position of deadwood, and the associated microclimatic differences, can strongly influence the community assembly after tree species identity has defined initial species pool. For example, Larsson <u>Ekström *et al*. (*2024*)</u> reported that standing and lying deadwood hosted different assemblages for both fungi and beetles, with standing deadwood supporting more beetle species and lying deadwood more fungal species. Likewise, Atrena *et al*. (2020) demonstrated that fungal richness was highest on lying coarse deadwood, while standing coarse deadwood supported lower fungal species richness, reinforcing the importance of deadwood position for fungal communities. Our result that felled and uprooted deadwood often resembled one another biologically is therefore plausible, as both substrates are relatively moist deadwood types that are likely similar in microclimatic conditions.

### Species confined to certain trees and types of dead wood

As expected, indicator species analyses showed that many fungal and beetle taxa were confined to tree species and preferred particular deadwood types, sometimes regardless of the tree species identity. For fungi, the indicator taxa on birch were mainly ascomycetous and mostly cryptic taxa, while relatively few fungal indicators were wood-decaying macrofungi, i.e. taxa with obvious sporocarps. This result suggests that indicator species analysis identifies taxa that are restricted to particular deadwood tree species and types, not necessarily those that are most functionally important, at least for decomposition processes. Some dominant wood-decaying basidiomycetes such as *Fomitopsis pinicola* colonise several tree species and deadwood types making them less likely to emerge as indicators of deadwood types. In contrast, many ascomycetes are more specialized, hence more strongly selective to differences in moisture or wood chemistry and therefore shown as indicator taxa, even if their contribution to decomposition is smaller (Ottosson *et al*., 2015; Purahong *et al*., 2018b; Brabcová *et al*., 2022).

As for beetles, standing deadwood was characterized by a relatively narrow set of indicator taxa., whereas lying deadwood (e.g., felled and uprooted trees) supported a different set of indicator species. The similarity in beetle community composition and indicator taxa between felled and uprooted deadwood types further indicates that once wood reaches the forest floor, humidity, temperature and other microclimatic variables become more important than the way the tree died. Also, the interaction networks may depend on macro- or microhabitat context (Chamberlain *et al*., 2014). An interesting example of context-dependent interaction between a host fungi and consumer beetle was shown between beetle *Cis punctulatus* and its host species *Trichaptum abietinum.* The fungi was an indicator of coniferous trees, both standing and lying, while the beetle was an indicator of lying conifer trees. This pattern suggests that the presence of the consumer beetle is not necessarily constrained by the presence of the host fungi but also the usability of the host within specific microhabitats (Schigel, 2012; Ryvarden & Melo, 2017). Overall, these results show that it is important to maintain a diversity of both deadwood tree species and deadwood types if the aim is to preserve not just species compositions, but also the functional and multitrophic associations linked to deadwood.

### Tree species and deadwood types explain the correlations between fungal and beetle communities

Our prediction that tree species and deadwood type explain fungal–beetle community multivariate covariation was only partly supported. The key point is that a significant RV coefficient does not demonstrate pairwise interactions between fungi and beetles; rather, it shows that the two communities change in parallel across samples more than expected by chance (Escoufier, 1973; Robert & Escoufier, 1976; Josse *et al*., 2008). Considering this explanation, the significant RV coefficient values in pine and spruce indicate a strong coupling between that fungal and beetle communities in conifer deadwood. In contrast, the non-significant RV coefficients in birch may suggest that such links were weaker or harder to detect potentially due to a smaller sample size. Because birch had fewer deadwood type combinations (i.e., burned standing and felled trees only) and a smaller sample size, reduced statistical power is a possible partial explanation; however, it is also biologically plausible that fungal and beetle communities are less correlated (e.g., have fewer links) compared to conifer trees. This could be partly explained by a high number of generalist beetle species in deciduous dead wood (Bergmark *et al*., 2024). Further, deciduous deadwood has faster decay rates than conifers (Mäkinen *et al*., 2006; Weedon *et al*., 2009) and according to Ramírez-Hernández *et al*. (2019) the interaction between late decay wood and associated beetles became weaker with advanced decay stages.

Burned standing spruce and pine were characterized by strong links between fungal and beetle communities. This is consistent with fire acting as an environmental filter: burning changes bark retention, wood surface conditions, and selecting for certain fungi and fire-favored beetle species (Hjältén *et al*., 2017; Fredriksson *et al*., 2025). For example, fire-scorching has been shown to reduce beetle richness, accelerate bark loss, and increase ascomycete fungi and insects feeding on them (Wikars, 2002). The strong community correlation in girdled spruce may reflect a different mechanism. Girdling is often slowly killing tree (Noel, 1970; Fujimori, 2001; Hägglund & Hjältén, 2018). Under these conditions, beetles may both track and facilitate fungal establishment (Zhao *et al*., 2019; Kandasamy *et al*., 2023), as insects in spruce can disperse fungal propagules and impact early fungal succession (Persson *et al*., 2011), leading to stronger covariation of fungal and beetle communities. As for the felled and uprooted deadwood types, the weaker covariation between fungal and beetle communities does not necessarily indicate random assembly, but rather a more heterogeneous environment in which fungi and beetles can occupy diverse and only partially overlapping niches (Parisi *et al*., 2018; Vítková *et al*., 2018). To sum up, these results suggest that fungal–beetle community correlations are the strongest where deadwood type conditions select for more specialized taxa, i.e., burned deadwood, but weaker where deadwood offers a range of environmental niches.

### Fungal-beetle network structure is modular and “anti-nested”, pattern consistent across tree species and tree species x deadwood types

Across three tree species and deadwood types, networks were consistently less nested and more modular than expected by chance. Thus, the fungal-beetle networks are not built around a few very common generalist species that share many partners. Instead, species tended to form groups of co-occurring species (aka modules), and the low nestedness suggests that there is little overlap among species sets, so many co-occurrences were relatively unique. In practice, this suggests low redundancy: if one fungus or beetle disappears, the links it has may not be easily replaced by another species. Interestingly, similar results were shown in the study by <u>Jacobsen *et al*. (2018)</u> in which it was demonstrated that deadwood fungal-beetle networks were anti-nested, moderately specialized and highly modular.

Contrary to our expectations, the strongest examples of high modularity and low nestedness were found for conifer deadwood species, especially felled spruce and uprooted pine. A likely explanation for this combination of high modularity and low nestedness is that deadwood is a highly patchy and temporary (ephemeral) habitat, and each deadwood unit develops its diverse microenvironmental conditions, including differences in moisture, temperature and bark cover (Zibold *et al*., 2024). Together with priority effects, fungal competition, and stochasticity, this leads to different fungal assemblages among deadwood units (Hiscox *et al*., 2018; Dahlberg *et al*., 2025). If beetles respond to the fungal niches rather than simply to fungal presence, species will form separate modules with little overlap, producing low nestedness and high modularity. The same process may explain why felled and uprooted deadwood showed weaker fungal–beetle community-level correlations but still strong modularity: lying wood provides heterogeneous conditions, including differences in moisture, bark loss, decay stage, and microclimate, that allow many different combinations of fungi and beetles to establish. This heterogeneity may be further increased by species that modify the substrate, such as wood-boring insects (Zuo *et al*., 2016; Sanders & Frago, 2024). This interpretation is consistent with the idea that habitat heterogeneity can increase niche diversity and promote compartmentalized network structure (Fujii *et al*., 2023). This also fits the species-level results, where most taxa were peripheral and confined to single modules, whereas connectors and hubs were rare. Connector taxa were more common in felled and uprooted wood, while standing deadwood, especially high stumps and girdled trees, was dominated by peripheral taxa, suggesting that the drier, more exposed conditions in standing wood limit colonization and lead to more isolated groups of co-occurring species (Hagge *et al*., 2024; Larsson Ekström *et al*., 2024a). Burned wood also had few connectors and hubs, consistent with fire acting as a strong filter on fungal and beetle communities.

## Conclusions

Our study shows that tree species was the main factor shaping both fungal and beetle communities, followed by deadwood type which impacted which species occurred within each tree species. The most evident differences in fungal and beetle community composition were between standing deadwood (girdled trees, high stumps, and often burned standing trees) and lying deadwood (felled and uprooted trees). A second (main) result was that fungal–beetle community correlations were not equally strong across all types of deadwood. They were strongest in conifers, especially in burned standing pine, burned standing spruce, and girdled spruce. This suggests that fungi and beetles become more closely linked when deadwood conditions strongly filter which species can colonize. In contrast, weaker community-level correlations in felled and uprooted wood likely reflect a more heterogenous environment. Importantly, our study showed that across all tree species and deadwood types, fungal–beetle networks were less nested and more modular than expected by chance. In simple terms, this means that the networks were not built around a few abundant generalist species. Instead, species formed small, separate groups of co-occurring taxa, with relatively little overlap among those groups. This suggests that many fungus–beetle links are specialized and may not be easily replaced if species are lost. The most modular networks were detected in conifer deadwood, especially felled spruce and uprooted pine. Future studies should follow these communities across later decay stages, test whether beetles respond mainly to fungal presence, fungal fruit bodies, or shared habitat conditions, and examine whether modular, low-nested networks differ in decomposition rate, nutrient turnover, or sensitivity to species loss. It will also be important to test which processes create these network modules, especially the roles of fungal competition, substrate heterogeneity, and microclimate.

