## Supporting Information for "Fungal–beetle networks in deadwood are modular and shaped by tree species and deadwood type"

**Methods S1:** Details on fungal DNA extraction, PCR amplification and sequencing steps

The sawdust samples used for DNA extraction were freeze-dried in their collected 50 mL Falcon tubes at -105 °C for 48 hours in a VirTis SP Scientific Freeze Dryer (SP Industries Inc., Suffolk, UK). For DNA extraction, 0.5 g of freeze-dried sawdust from each sample was placed separately into a 2 mL screw cap centrifugation tube. DNA was extracted with the Techtum NucleoSpin soil kit (Macherey-Nagel, Düren, Germany) according to manufacturer’s recommendations. For extractions, was used SL1 lysis buffer with enhancer Sx. The quality and quantity of resulting DNA products was evaluated by an ND-1000 spectrophotometer (NanoDrop Technologies, Wilmington, DE, USA). The ITS2 region was amplified using the fungal-specific primers gITS7 and a 3:1 mix of the reverse primers ITS4 and ITS4arch, with both forward and reverse primers fitted with unique 8 bp sample identification tags resulting in amplicons 250-400bp in length (Ihrmark *et al.*, 2012; Clemmensen *et al.*, 2016). DNA extracts were performed in 50-μl PCR reaction containing 5 µL of DNA template volume and amplified using the following cycling programme: 5 min at 95°C; 20-35cycles for 30 s at 95°C, 30 s at 56°C and 30 s at 72°C, followed by a final elongation step for 7 min at 72°C. The number of PCR cycles was optimized for each sample by re-running samples with too strong or too weak PCR products (according to gel electrophoresis) with cycle numbers adopted to obtain “weak but visible” bands on the gel (Castaño et al. 2020). PCR products were run in duplicates, which were pooled and cleaned using the AMPure kit (Beckman Coulter Inc., Brea, CA, USA). DNA concentrations were established using a Qubit fluorometer (Life Technologies, Carlsbad, CA, USA), and DNA from each sample were pooled together and final pool was purified again using the E.Z.N.A. Omega cycle pure kit (Omega Bio-tek, Norcross, GA, USA). Amplicon size distribution was pre-checked using the 2100 Bioanalyzer system (Agilent, Santa Clara, CA, USA) and composite samples were sequenced on the RSII platforms (Pacific Biosciences, Menlo Park, CA, USA) by SciLifeLab NGI (Uppsala, Sweden) after addition of sequencing adaptors by ligation. The PacBio platform was chosen to minimize bias due to size variation in the amplicon pool, which is considerable for the fungal ITS2 region (Castaño *et al.*, 2020).

Sequences were filtered and clustered using the SCATA pipeline (scata.mykopat.slu.se; (Ihrmark *et al.*, 2012). Sequences were quality checked to remove sequences shorter than 100 bp, with mean quality scores lower than 20, with individual bases with a quality score lower than 3, or with a missing 3’ or 5’ tag. Sequences were screened for the gITS7 and ITS4 primers, requiring a minimum match of 90%, and reverse complemented if necessary. Globally unique genotypes were removed to reduce the incidence of sequencing errors. Quality filtering removed 39% of the total sequences, and another 18% were removed as unique genotypes. Remaining sequences were clustered into species hypotheses (hereafter species; (Kõljalg *et al.*, 2013) through pairwise comparisons with USEARCH (Edgar, 2010) followed by single linkage clustering, with the minimum similarity to the closest neighbour required to enter a cluster set at 98.5%. To reduce the impact of long deletions, gap elongation penalty was reduced during clustering. Homopolymers were collapsed to 3 bp. Non-fungal sequence clusters were identified and removed by blast comparisons with the NCBI nt database. Fungal species were identified by blast comparisons with the UNITE database (Abarenkov *et al.*, 2024).

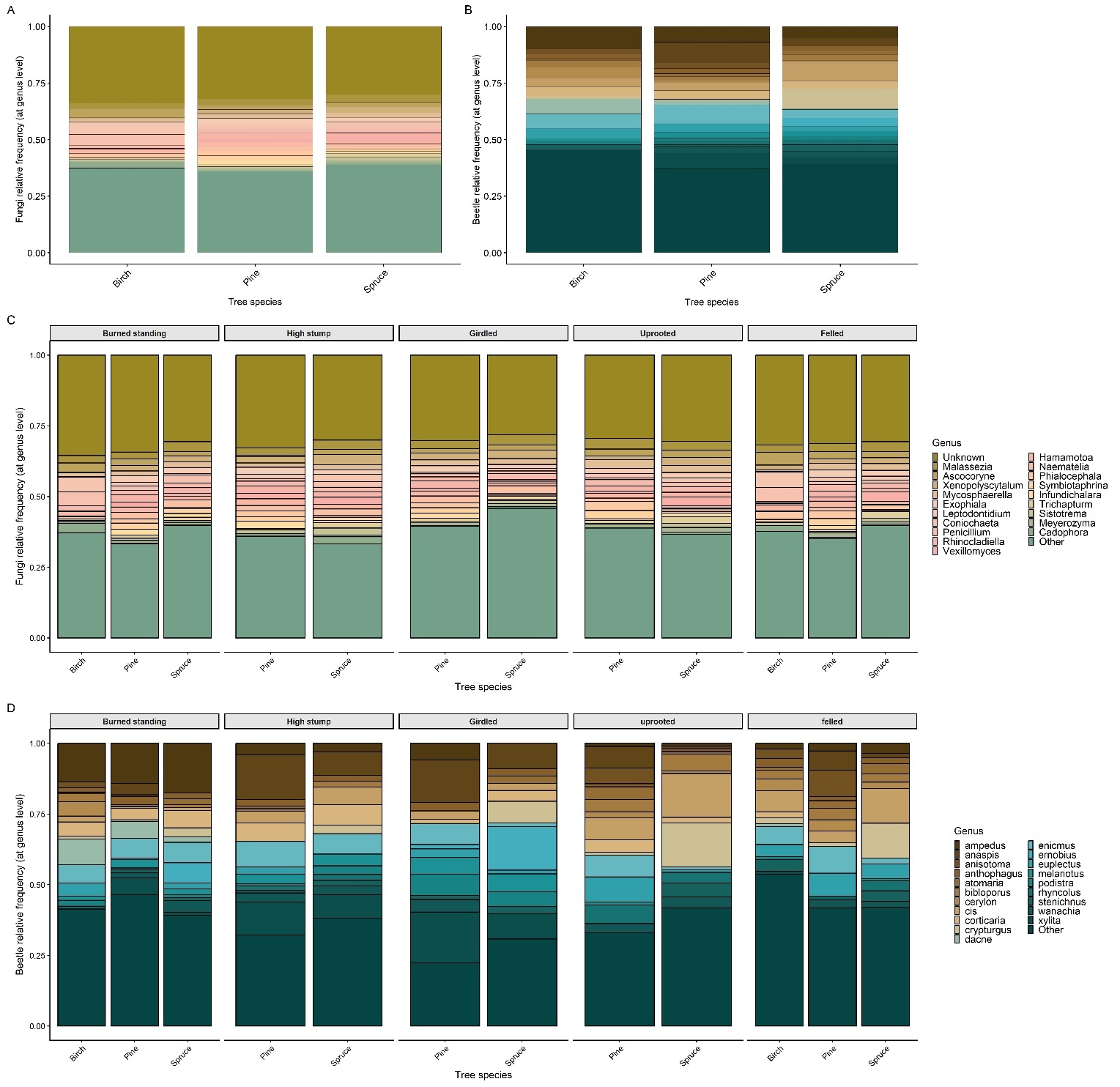

**Figure S1.** *Relative frequencies of fungal and beetle genera across tree species and deadwood types*.
Stacked bar charts show the relative genus-level frequencies of fungi and beetles across three tree species (Panels A-B) and five deadwood types (Panels C-D). For fungi, relative frequency was calculated as the proportion of samples in which each genus was detected based on presence/absence data from sequencing reads. For beetles, relative frequency represents the proportion of samples where individuals from each genus were recorded.
The 20 most frequently observed fungal and beetle genera are shown; all remaining genera are grouped as "Other".

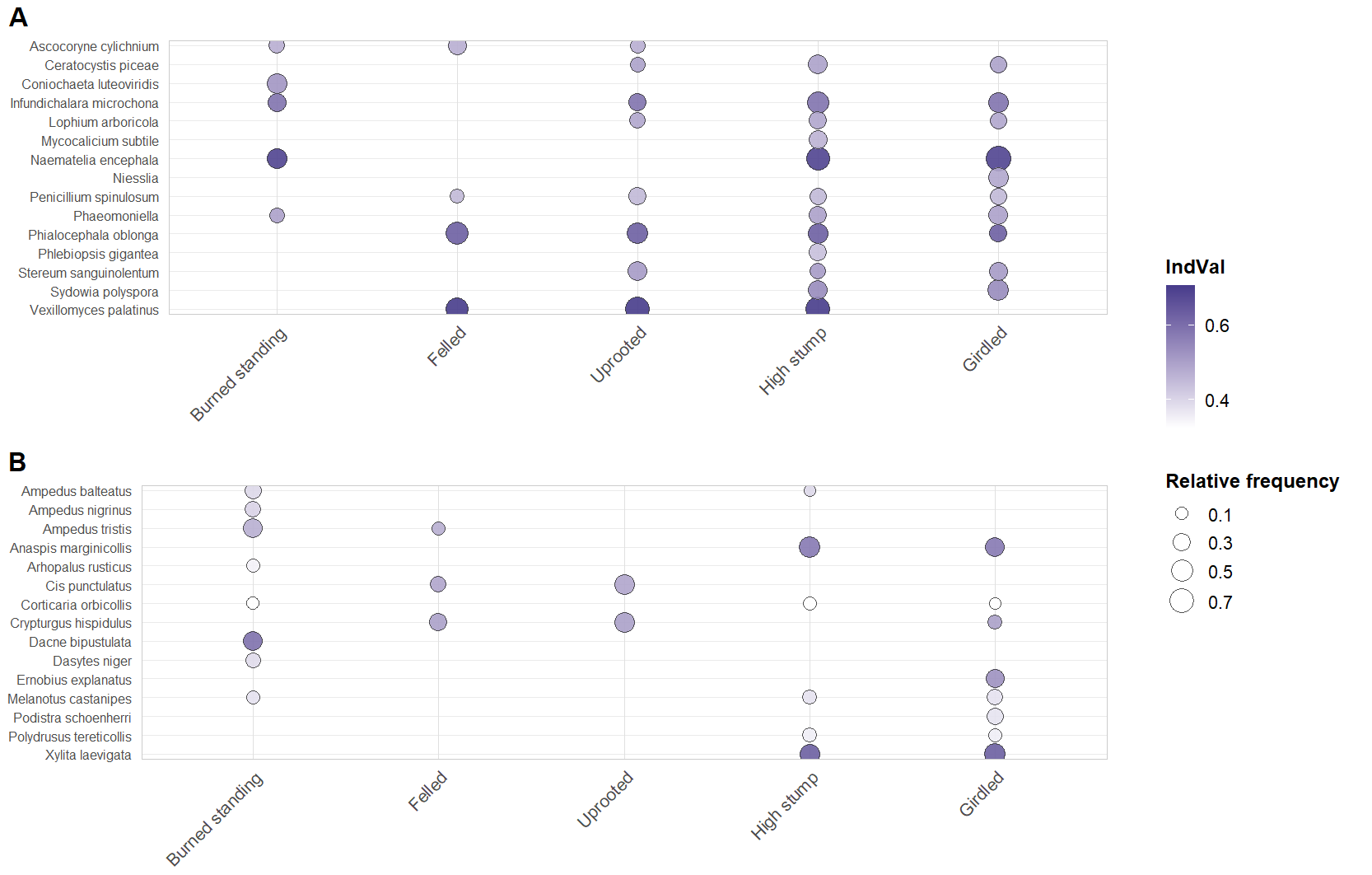

**Figure S2*.*** *Indicator taxa across deadwood types. Bubble plots show taxa identified as significant indicators for (A) fungi and (B) beetles.* Bubble color represents indicator value (IndVal; darker violet = stronger association), and bubble size represents the taxon’s relative frequency (i.e., proportion of samples in which the taxon was detected) within each group. For each group, taxa ranked by IndVal and ordered alphabetically on the y-axis.

**Table S1**. Coordinates of the gap-cut and burned stands.

| **Stand**  **coordinates**  **Stand ID** | **Latitude** | **Longitude** |
| --- | --- | --- |
| Burned 1935 | 17.4727460006673 | 63.3950799977972 |
| Burned 2746 | 20.0971819999584 | 64.5019240000445 |
| Burned 3126 | 17.4557370002981 | 63.4068720040526 |
| Burned 4402 | 17.4854860004218 | 64.3151840015354 |
| Burned 6210 | 18.2862699987325 | 63.9568069975949 |
| Burned 7552 | 18.7950810000006 | 64.0452580000381 |
| Gap 505 | 17.6222460005144 | 63.4685949987676 |
| Gap 4848 | 18.3635919998419 | 63.9414779963013 |
| Gap 5655 | 18.7983130003202 | 64.0273250021821 |
| Gap 6323 | 20.2104168202451 | 64.259771809051 |
| Gap 7315 | 18.3014700000127 | 63.9653950000363 |
| Gap 8570 | 17.6315000000077 | 64.3497540000334 |

**Table S2**. Summary of number of replicates sampled for beetle and fungal community analyses across deadwood types (burned, felled, high stump, girdled, uprooted) and tree species (spruce, pine, birch) in burned and gap stands. Each study site was initially designed to include five replicates for every deadwood type–tree species combination. One sample corresponds to one tree: for beetles, a single emergence trap was placed on each tree; for fungi, three drill-hole cores were taken at the trap location and pooled into one composite sample. Some replicates were lost due to sampling or processing issues, such as failed beetle trap collections or unsuccessful fungal DNA amplification. The final datasets comprised 348 beetle samples and 356 fungal samples. For the locations of the stands see Figure 1 and Table S1.

|  | Burned | | | Felled | | | High stump | | Girdled | | Uprooted | | |
| --- | --- | --- | --- | --- | --- | --- | --- | --- | --- | --- | --- | --- | --- |
|  | *Spruce* | *Pine* | *Birch* | *Spruce* | *Pine* | *Birch* | *Spruce* | *Pine* | *Spruce* | *Pine* | | *Spruce* | *Pine* |
| *A. Beetles* |  |  |  |  |  |  |  |  |  |  | |  |  |
| Burned 1935 | 5 | 5 | 5 |  |  |  |  |  |  |  | |  |  |
| Burned 2746 | 4 | 5 | 5 |  |  |  |  |  |  |  | |  |  |
| Burned 3126 | 4 | 5 | 4 |  |  |  |  |  |  |  | |  |  |
| Burned 4402 | 5 | 5 | 4 |  |  |  |  |  |  |  | |  |  |
| Burned 6210 | 5 | 5 | 4 |  |  |  |  |  |  |  | |  |  |
| Burned 7552 | 5 | 5 | 5 |  |  |  |  |  |  |  | |  |  |
| Gap 505 |  |  |  | 5 | 4 | 4 | 4 | 4 | 5 | 5 | | 5 | 4 |
| Gap 4848 |  |  |  | 4 | 2 | 5 | 5 | 5 | 4 | na | | 5 | 5 |
| Gap 5655 |  |  |  | 5 | 3 | 4 | 5 | 4 | 5 | 1 | | 5 | na |
| Gap 6323 |  |  |  | 5 | 4 | 4 | 5 | 5 | 3 | 4 | | 5 | 4 |
| Gap 7315 |  |  |  | 5 | 2 | 3 | 4 | 3 | 4 | 4 | | 4 | 3 |
| Gap 8570 |  |  |  | 5 | 5 | 5 | 3 | 4 | 1 | 3 | | 4 | na |
| *B. Fungi* |  |  |  |  |  |  |  |  |  |  | |  |  |
| Burned 1935 | 5 | 5 | 5 |  |  |  |  |  |  |  | |  |  |
| Burned 2746 | 5 | 5 | 4 |  |  |  |  |  |  |  | |  |  |
| Burned 3126 | 5 | 5 | 5 |  |  |  |  |  |  |  | |  |  |
| Burned 4402 | 5 | 5 | 5 |  |  |  |  |  |  |  | |  |  |
| Burned 6210 | 5 | 5 | 5 |  |  |  |  |  |  |  | |  |  |
| Burned 7552 | 5 | 5 | 5 |  |  |  |  |  |  |  | |  |  |
| Gap 505 |  |  |  | 5 | 4 | 3 | 5 | 5 | 5 | 4 | | 3 | 4 |
| Gap 4848 |  |  |  | 5 | 2 | 5 | 5 | 5 | 4 | na | | 5 | 5 |
| Gap 5655 |  |  |  | 5 | 2 | 4 | 5 | 5 | 4 | 1 | | 5 | 2 |
| Gap 6323 |  |  |  | 4 | 4 | 4 | 4 | 3 | 5 | 4 | | 4 | 3 |
| Gap 7315 |  |  |  | 5 | 3 | 3 | 5 | 4 | 5 | 5 | | 5 | 3 |
| Gap 8570 |  |  |  | 5 | 5 | 4 | 3 | 4 | 2 | 2 | | 5 | 1 |

**Table S3.** Species roles in fungal-beetle networks across tree species. Species’ structural roles were quantified using within-module degree (z) and participation coefficient (c). Species were classified as peripherals (low z, low c), connectors (low z, high c), module hubs (high z, low c), or network hubs (high z, high c). The table reports the top-ranked taxa per category (tree species × taxon group × role) based on role-specific z/c ranking.

| Tree species | Taxon group | Network role | Taxon | Within-module connectivity (z) | Among-module connectivity (c) | Ranking score |
| --- | --- | --- | --- | --- | --- | --- |
| Birch | Beetles | Peripheral | *Anisotoma glabra* | 1,65 | 0,4 | -0,88 |
| Birch | Beetles | Peripheral | *Latridius hirtus* | 1,17 | 0,56 | -1,34 |
| Birch | Beetles | Peripheral | *Stenotrachelus aeneus* | 0,56 | 0,27 | -1,97 |
| Birch | Beetles | Peripheral | *Cis boleti* | 0,29 | 0,6 | -2,21 |
| Birch | Beetles | Peripheral | *Athous subfuscus* | 0,32 | 0,1 | -2,24 |
| Birch | Beetles | Peripheral | *Triplax russica* | -0,29 | 0,47 | -2,79 |
| Birch | Beetles | Peripheral | *Schizotus pectinicornis* | -0,39 | 0,59 | -2,89 |
| Birch | Beetles | Peripheral | *Coryphium angusticolle* | -0,41 | 0,39 | -2,92 |
| Birch | Beetles | Peripheral | *Anisotoma castanea* | -0,56 | 0,45 | -3,06 |
| Birch | Beetles | Peripheral | *Hylobius abietis* | -0,61 | 0,58 | -3,11 |
| Birch | Beetles | Connector | *Ampedus tristis* | 0,59 | 0,74 | 0,74 |
| Birch | Beetles | Connector | *Dacne bipustulata* | 0,95 | 0,74 | 0,74 |
| Birch | Beetles | Connector | *Elateroides dermestoides* | -1,77 | 0,74 | 0,74 |
| Birch | Beetles | Connector | *Enicmus rugosus* | 1,06 | 0,74 | 0,74 |
| Birch | Beetles | Connector | *Bibloporus minutus* | 0,59 | 0,74 | 0,74 |
| Birch | Beetles | Connector | *Corticaria orbicollis* | -0,47 | 0,74 | 0,74 |
| Birch | Beetles | Connector | *Enicmus fungicola* | 0 | 0,73 | 0,73 |
| Birch | Beetles | Connector | *Orchesia micans* | -0,95 | 0,73 | 0,73 |
| Birch | Beetles | Connector | *Rhizophagus dispar* | -1,24 | 0,73 | 0,73 |
| Birch | Beetles | Connector | *Scaphisoma agaricinum* | 1,15 | 0,73 | 0,73 |
| Birch | Fungi | Peripheral | *Helicogloea pellucida* | 1,54 | 0,59 | -0,96 |
| Birch | Fungi | Peripheral | *Peniophorella praetermissa* | 1,24 | 0,54 | -1,27 |
| Birch | Fungi | Peripheral | *Mollisia cinerea* | 1,24 | 0,4 | -1,28 |
| Birch | Fungi | Peripheral | *Exidia candida* | 1,21 | 0,58 | -1,29 |
| Birch | Fungi | Peripheral | *Subulicystidium longisporum* | 0,88 | 0,56 | -1,62 |
| Birch | Fungi | Peripheral | *Fomitopsis betulina* | 0,64 | 0,44 | -1,87 |
| Birch | Fungi | Peripheral | *Solicoccozyma terricola* | 0,64 | 0,41 | -1,87 |
| Birch | Fungi | Peripheral | *Penicillium paxilli* | 0,22 | 0,3 | -2,3 |
| Birch | Fungi | Peripheral | *Scheffersomyces lignosus* | 0,11 | 0,49 | -2,4 |
| Birch | Fungi | Peripheral | *Coniochaeta luteorubra* | -0,11 | 0,52 | -2,61 |
| Birch | Fungi | Connector | *Phellinus ignarius* | 0,3 | 0,75 | 0,75 |
| Birch | Fungi | Connector | *Cadophora malorum* | 2,18 | 0,74 | 0,74 |
| Birch | Fungi | Connector | *Phialocephala oblonga* | 2,18 | 0,74 | 0,74 |
| Birch | Fungi | Connector | *Ascocoryne cylichnium* | 2,18 | 0,74 | 0,74 |
| Birch | Fungi | Connector | *Coniochaeta* | 1,51 | 0,74 | 0,74 |
| Birch | Fungi | Connector | *Leptodontidium trabinellum* | 1,51 | 0,74 | 0,74 |
| Birch | Fungi | Connector | *Hamamotoa lignophila* | 1,71 | 0,74 | 0,74 |
| Birch | Fungi | Connector | *Phellinus nigricans* | -1,11 | 0,74 | 0,74 |
| Birch | Fungi | Connector | *Hyphoderma setigerum* | 0,77 | 0,74 | 0,74 |
| Pine | Beetles | Peripheral | *Anisotoma castanea* | 2,33 | 0,52 | -0,19 |
| Pine | Beetles | Peripheral | *Cerylon histeroides* | 1,86 | 0,27 | -0,73 |
| Pine | Beetles | Peripheral | *Ptinella johnsoni* | 1,13 | 0,57 | -1,37 |
| Pine | Beetles | Peripheral | *Gabrius expectatus* | 1,09 | 0,45 | -1,42 |
| Pine | Beetles | Peripheral | *Aspidiphorus orbiculatus* | 1,05 | 0,52 | -1,46 |
| Pine | Beetles | Peripheral | *Euplectus punctatus* | 0,66 | 0,6 | -1,84 |
| Pine | Beetles | Peripheral | *Atomaria bella* | 0,57 | 0,53 | -1,93 |
| Pine | Beetles | Peripheral | *Anaspis rufilabris* | 0,23 | 0,59 | -2,27 |
| Pine | Beetles | Peripheral | *Rhyncolus ater* | 0,23 | 0,46 | -2,27 |
| Pine | Beetles | Peripheral | *Anastrangalia reyi* | 0,14 | 0,61 | -2,36 |
| Pine | Beetles | Connector | *Enicmus rugosus* | 1,74 | 0,74 | 0,74 |
| Pine | Beetles | Connector | *Corticaria rubripes* | 0,19 | 0,73 | 0,73 |
| Pine | Beetles | Connector | *Corticarina parvula* | -0,78 | 0,73 | 0,73 |
| Pine | Beetles | Connector | *Anthophagus omalinus* | 0,58 | 0,73 | 0,73 |
| Pine | Beetles | Connector | *Xylita laevigata* | 1,9 | 0,72 | 0,72 |
| Pine | Beetles | Connector | *Bibloporus minutus* | -0,05 | 0,72 | 0,72 |
| Pine | Beetles | Connector | *Wanachia triguttata* | 0,82 | 0,72 | 0,72 |
| Pine | Beetles | Connector | *Anaspis marginicollis* | 2,22 | 0,71 | 0,71 |
| Pine | Beetles | Connector | *Melanotus castanipes* | 0,47 | 0,7 | 0,7 |
| Pine | Beetles | Connector | *Coryphium angusticolle* | -0,19 | 0,7 | 0,7 |
| Pine | Fungi | Peripheral | *Sarea difformis* | 2,31 | 0,57 | -0,2 |
| Pine | Fungi | Peripheral | *Cosmospora arxii* | 1,44 | 0,6 | -1,06 |
| Pine | Fungi | Peripheral | *Fomes fomentarius* | 1,15 | 0,47 | -1,36 |
| Pine | Fungi | Peripheral | *Mortierella humilis* | 1,08 | 0,57 | -1,43 |
| Pine | Fungi | Peripheral | *Syzygospora effibulata* | 1,08 | 0,57 | -1,43 |
| Pine | Fungi | Peripheral | *Cortinarius causticus* | 1,08 | 0,46 | -1,43 |
| Pine | Fungi | Peripheral | *Hyaloscypha variabilis* | 1,08 | 0,46 | -1,43 |
| Pine | Fungi | Peripheral | *Aspergillus* | 0,86 | 0,51 | -1,65 |
| Pine | Fungi | Peripheral | *Hypochnicium albostramineum* | 0,79 | 0,62 | -1,71 |
| Pine | Fungi | Peripheral | *Xylopsora* | 0,79 | 0,56 | -1,71 |
| Pine | Fungi | Connector | *Orbilia* | -1,11 | 0,77 | 0,77 |
| Pine | Fungi | Connector | *Piskurozyma* | -1,11 | 0,77 | 0,77 |
| Pine | Fungi | Connector | *Xylodon novozelandicus* | -0,87 | 0,76 | 0,76 |
| Pine | Fungi | Connector | *Clonostachys rosea* | -0,22 | 0,75 | 0,75 |
| Pine | Fungi | Connector | *Phlebiopsis gigantea* | -0,57 | 0,75 | 0,75 |
| Pine | Fungi | Connector | *Rhodotorula mucilaginosa* | 1,08 | 0,75 | 0,75 |
| Pine | Fungi | Connector | *Oidiodendron pilicola* | 1,08 | 0,74 | 0,74 |
| Pine | Fungi | Connector | *Naganishia adeliensis* | 1,08 | 0,74 | 0,74 |
| Pine | Fungi | Connector | *Coniochaeta* | 2,26 | 0,74 | 0,74 |
| Pine | Fungi | Connector | *Sporobolomyces roseus* | -0,55 | 0,74 | 0,74 |
| Spruce | Beetles | Peripheral | *Xylita laevigata* | 2,26 | 0,62 | -0,24 |
| Spruce | Beetles | Peripheral | *Sepedophilus immaculatus* | 1,53 | 0,56 | -0,97 |
| Spruce | Beetles | Peripheral | *Acrulia inflata* | 1,09 | 0,49 | -1,42 |
| Spruce | Beetles | Peripheral | *Athous subfuscus* | 0,63 | 0,53 | -1,88 |
| Spruce | Beetles | Peripheral | *Pteryx suturalis* | 0,43 | 0,52 | -2,07 |
| Spruce | Beetles | Peripheral | *Cerylon histeroides* | -0,18 | 0,61 | -2,68 |
| Spruce | Beetles | Peripheral | *Ampedus balteatus* | -0,33 | 0,59 | -2,83 |
| Spruce | Beetles | Peripheral | *Agathidium seminulum* | -0,5 | 0,59 | -3 |
| Spruce | Beetles | Peripheral | *Euplectus karstenii* | -0,5 | 0,58 | -3 |
| Spruce | Beetles | Peripheral | *Stephanopachys substriatus* | -0,62 | 0,6 | -3,12 |
| Spruce | Beetles | Connector | *Stenichnus bicolor* | -0,33 | 0,8 | 0,8 |
| Spruce | Beetles | Connector | *Cis punctulatus* | 1,23 | 0,79 | 0,79 |
| Spruce | Beetles | Connector | *Ampedus tristis* | -0,3 | 0,79 | 0,79 |
| Spruce | Beetles | Connector | *Crypturgus hispidulus pusillus* | 1,14 | 0,79 | 0,79 |
| Spruce | Beetles | Connector | *Dryocoetes autographus* | 2,21 | 0,78 | 0,78 |
| Spruce | Beetles | Connector | *Orthoperus rogeri* | 0,54 | 0,78 | 0,78 |
| Spruce | Beetles | Connector | *Bibloporus minutus* | 1,78 | 0,78 | 0,78 |
| Spruce | Beetles | Connector | *Wanachia triguttata* | -0,36 | 0,77 | 0,77 |
| Spruce | Beetles | Connector | *Malthodes guttifer* | -0,51 | 0,77 | 0,77 |
| Spruce | Beetles | Connector | *Hadreule elongatula* | -1,08 | 0,77 | 0,77 |
| Spruce | Fungi | Peripheral | *Niesslia* | 2,44 | 0,59 | -0,06 |
| Spruce | Fungi | Peripheral | *Pichia holstii* | 2,24 | 0,35 | -0,38 |
| Spruce | Fungi | Peripheral | *Cosmospora arxii* | 2,04 | 0,55 | -0,46 |
| Spruce | Fungi | Peripheral | *Coniophora olivacea* | 1,51 | 0,6 | -0,99 |
| Spruce | Fungi | Peripheral | *Carcinomyces polyporinus* | 1,51 | 0,54 | -1 |
| Spruce | Fungi | Peripheral | *Cytospora* | 1,44 | 0,52 | -1,06 |
| Spruce | Fungi | Peripheral | *Exidia saccharina* | 1,24 | 0,48 | -1,27 |
| Spruce | Fungi | Peripheral | *Oidiodendron pilicola* | 1,12 | 0,61 | -1,38 |
| Spruce | Fungi | Peripheral | *Mycocalicium subtile* | 1,04 | 0,61 | -1,46 |
| Spruce | Fungi | Peripheral | *Phaeomoniella* | 1,04 | 0,55 | -1,46 |
| Spruce | Fungi | Connector | *Debaryomyces hansenii* | -0,51 | 0,8 | 0,8 |
| Spruce | Fungi | Connector | *Aureobasidium proteae* | 0,68 | 0,79 | 0,79 |
| Spruce | Fungi | Connector | *Neoantrodia serialis* | 0,25 | 0,79 | 0,79 |
| Spruce | Fungi | Connector | *Oidiodendron griseum* | 0,62 | 0,78 | 0,78 |
| Spruce | Fungi | Connector | *Vishniacozyma victoriae* | -0,2 | 0,78 | 0,78 |
| Spruce | Fungi | Connector | *Pezoloma ericae* | -1,31 | 0,78 | 0,78 |
| Spruce | Fungi | Connector | *Gloeophyllum sepiarium* | 0,68 | 0,78 | 0,78 |
| Spruce | Fungi | Connector | *Peniophorella praetermissa* | 0,69 | 0,78 | 0,78 |
| Spruce | Fungi | Connector | *Dacromyces stillatus* | 0,47 | 0,78 | 0,78 |
| Spruce | Fungi | Connector | *Calocera furcata* | 1,36 | 0,78 | 0,78 |
| Spruce | Fungi | Module hub | *Sydowia polyspora* | 2,84 | 0,51 | 2,84 |
| Spruce | Fungi | Module hub | *Niesslia tenuis* | 2,64 | 0,51 | 2,64 |
| Spruce | Fungi | Network hub | *Naematelia encephala* | 3,04 | 0,71 | 3,76 |
| Spruce | Fungi | Network hub | *Serpula himantioides* | 2,51 | 0,74 | 3,26 |
